# The HOMEODOMAIN-Like protein HDL mediates chromatin organization and rewires leaf epidermal patterning

**DOI:** 10.1101/2023.09.13.557516

**Authors:** Ansar Ali, Chi Kuan, Fu-Yu Hung, Tsai-Chen Chen, Hui-Chun Lee, Shao-Li Yang, Yun-Ru Feng, Keqiang Wu, Chin-Min Kimmy Ho

## Abstract

The *Arabidopsis* leaf epidermis is comprised of trichomes, pavement cells, and stomata, originating from versatile precursor cells capable of dividing or differentiating to create unique epidermal patterns. The mechanism governing these transitions and the maintenance of cell heterogeneity remains unclear. In this study, we identified a novel <u>h</u>omeo<u>d</u>omain-<u>l</u>ike superfamily protein, HDL, localized in chromocenters and playing a role in chromatin organization. HDL interacts with <u>h</u>istone <u>d</u>e<u>a</u>cetylase 6 (HDA6) and methyltransferases, indicating its function in modulating chromatin accessibility. In *hdl* mutants, we observed moderately increased chromatin accessibility in the promoter region of protein-coding genes, along with reduced stomatal density and elevated trichome numbers on leaf surfaces. Corresponding to these phenotypes, stomatal-related gene expression decreased, while a transcriptional reporter for *GLABRA2*, a trichome initiation gene, exhibited higher and more variable expression levels. These findings highlight how HDL-mediated chromatin organization influences epidermal cell fate by modulating gene expression and enhancing cell heterogeneity within the leaf epidermis.

**One Sentence Summary:** A homeodomain-like protein functions with histone modifiers to regulate chromatin and influence cell fate decisions.

---

Chromatin is the complex of DNA and proteins that make up the chromosomes within the nucleus of eukaryotic cells and plays a central role in gene regulation and genome organization. Histones with specific post-translational modifications and various chromatin-associated proteins contribute to chromatin dynamics by influencing access to DNA regulatory regions and mediating different transcriptional outcomes (*1, 2*). Different histone modifications are associated with transcriptional activation or repression. Particularly di-methylation of histone 3 lysine 9 (H3K9me2) is one of the critical repression marks that correlates with heterochromatin formation (*3*).

Several epigenetic mechanisms are known to be important for regulation of cell division versus differentiation that is critical for meristem maintenance and development. The homeodomain transcription factor (TF), WUSCHEL (WUS) is an essential regulator of meristem maintenance (*4*). KNUCKLES (KNU) serves as a repressor of *WUS* and its expression is in turn suppressed by Polycomb group (PcG) proteins in the shoot apical meristem. During the initiation of floral development, the floral homeotic protein AGAMOUS triggers activation of *KNU* by modifying histones located at the *KNU* locus. This activation, in turn, results in the repression of *WUS* and subsequent flower development (*5, 6*). This example illustrates how epigenetic regulation is incorporated into a coordinated transcriptional program to control meristem maintenance. However, how chromatin structure impacts cell fate after cells cease cell division activity remains elusive. <u>S</u>tomatal lineage <u>g</u>round <u>c</u>ells (SLGCs) function as epidermal stem cells that can proliferate to generate more asymmetrically divided stomatal lineage cells or differentiate to become polyploid pavement cells or trichomes (Fig. 1A). Therefore, analyzing genes enriched in SLGCs can provide clues about the molecular mechanism underlying the cell state transition in developing leaves. Here, from our published SLGC gene list (*7*), we identify a previously uncharacterized <u>h</u>omeo<u>d</u>omain-<u>l</u>ike (HDL) protein coded by *AT3G53440* as an essential factor for chromatin organization and show that the loss of chromatin compaction favors cell differentiation rather than division, thus leading to the alternation of epidermal patterns.

**Figure 1.**
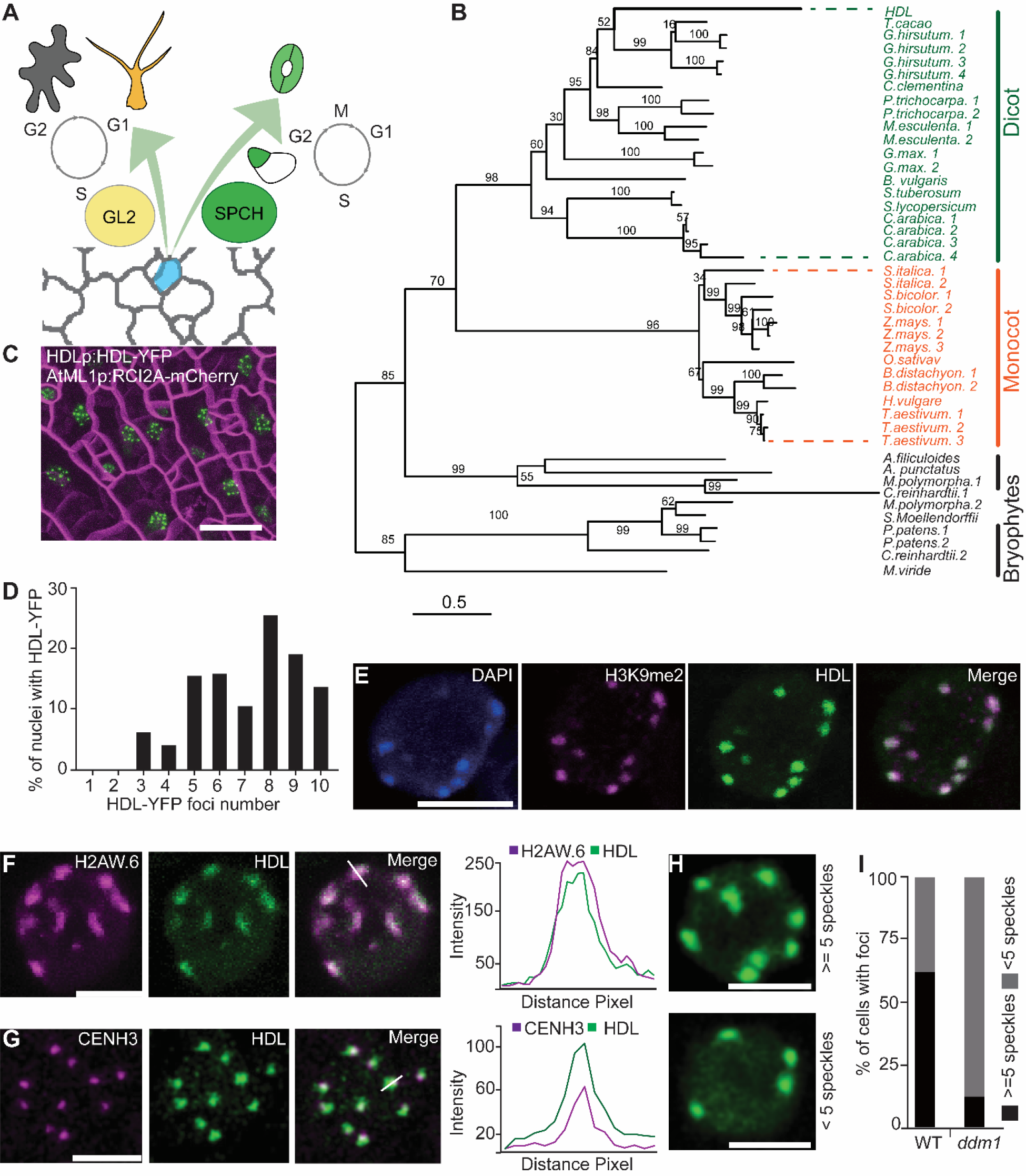
HDL associates with chromocenters. **(A)** The three distinct cell types of the leaf epidermis are produced through either cell division or differentiation. Protodermal cells (PD, blue) give rise to trichomes (yellow), stomata (green), and pavement cells (grey). The transcription factor (TF), GLABRA2 (GL2), drives PD cells to become polyploid trichome cells during trichome formation. SPEECHLESS (SPCH), the primary transcription factor governing the stomatal lineage, triggers an asymmetric cell division that yields a small meristemoid (M, green) and a stomatal lineage ground cell (SLGC, white), ultimately leading to the formation of a pair of guard cells. **(B)** Phylogenetic analysis of HDL homologs in different plant species indicates HDL is a green lineage-specific and single-copy gene in *Arabidopsis*. The number indicates the bootstrap value. **(C)** Translational fusion of HDL forms nuclear speckles (green) on leaf epidermis. Plasma membrane (PM) is visualized with *ML1p: RCI2A-mChery* (magenta). **(D)** Quantification of the nuclei with HDL nuclear speckles in 5-dpg (days post germination) abaxial true leaves (n=79 nuclei). **(E)** Immunostaining of root nuclei isolated from transgenic plants expressing *HDLp:HDL-YFP*, co-labeled with the heterochromatin marker H3K9me2. DNA counterstained with DAPI (Blue). **(F)** Confocal image of a root cell expressing *HDLp:HDL-YFP* and *H2AW*.*6p:H2AW*.*6-CFP*. **(G)** Confocal image of a leaf epidermal cell expressing *CENH3p:CENH3-GFP* and *HDLp:HDL-YFP*. The line graph represents the localization patterns of H2AW.6, CENH3, and HDL. **(H)** Representative images of HDL-YFP foci in nuclei as condensed (≥5 nuclear speckles) or decondensed (less than 5 nuclear speckles) in WT or *ddm1* plants. **(I)** Quantification of the proportion of nuclei exhibiting HDL-YFP foci in the wild type (WT) and *ddm1* mutant, n= 100 and 112 respectively. Scale bars, 20 µm in C, and 5 µm in E, F, G, and H.

## HDL associates with chromocenters

By analyzing overexpression lines of SLGC candidate genes, we found that plants expressing *35S:HDL-YFP* have more small cells on the leaf epidermis, indicating increased cell division of SLGCs (fig. S1). The phylogenetic analysis shows that HDL is a plant lineage-specific protein and is present as a single copy in *Arabidopsis* (Fig. 1B). The homeodomain of HDL is also predicted as a SANT/Myb-like DNA-binding domain which is present in the subunits of many chromatin-remodeling complexes in plants and animals (fig. S2A, Data S1) (*8, 9*). Endogenous promoter-driven HDL (*HDLp:HDL-YFP*) localized in nuclear speckles in leaf epidermis and root cells (Fig. 1C and fig. S2B). There were ten or less YFP speckles per cell, with a median of seven per nucleus (Fig. 1D). This observation resembles *Arabidopsis* heterochromatic chromocenters, where centromeric and pericentromeric regions of all five chromosomes form similar nuclear speckles when stained with DAPI (*10, 11*). To assess whether HDL is localized at heterochromatin chromocenters, we performed immunostaining of HDL-YFP nuclei with H3K9me2, a heterochromatin chromocenter marker. The results showed that HDL and H3K9me2 colocalized with chromocenters marked by dense DAPI staining (Fig. 1E and fig. S2, C and D). Additionally, HDL colocalized with another heterochromatin marker, H2AW.6, which marks constitutive heterochromatin and is required for chromatin condensation (*12, 13*) (Fig. 1F). When *HDLp:HDL-YFP* was crossed with *CENH3p:CENH3-GFP*, a centromere localized histone H3 variant, we found that HDL and CENH3 partially colocalized but HDL has a broader localization range (Fig. 1G), indicating that HDL occupies wide-range of chromatin domains.

Previously, it has been shown that DNA methylation is required for heterochromatin formation and that loss of the chromatin remodeler protein DECREASED DNA METHYLATION-1 (DDM1) disrupted chromocenters structure (*14, 15*). Therefore, to investigate whether the localization of HDL depends on chromocenters, we expressed *HDLp:HDL-YFP* in *ddm1* mutants. These plants had a reduced number of condensed chromocenters marked by HDL-YFP (90% of nuclei had less than five HDL-YFP speckles in *ddm1* vs. only 40% in WT) (Fig. 1, H and I). These results suggest that HDL associates with heterochromatin chromocenters.

## HDL is required for chromocenter formation and modulates developmental choices

To understand the function of HDL, a knock-down T-DNA insertional mutant, *hdl*, was primarily used in this study (fig. S3, A to F). We also generated a 481 bp *hdl-crispr* mutant using CRISPR-Cas9 editing (fig. S3, G to J). The *hdl* mutants had a clear phenotype of smaller leaves and reduced root length that could be complemented by *HDLp:HDL-YFP* (fig. S3, D to F). It has been reported that mutations in heterochromatin-associated proteins led to chromatin de-condensation (*13, 15, 16*). To test the hypothesis that HDL is localized in heterochromatic chromocenters and is important for heterochromatin formation, we used the T-DNA insertional mutant, *hdl*, to perform experiments in the following sections. Consistent with our hypothesis, DAPI staining found that the *hdl* had a reduced number of chromocenters (5-7 in *hdl* vs. 8-10 in WT); however, the size of each chromocenter was increased (Fig. 2, A and B and fig. S3, K and L). In addition, immunostaining with H3K9me2 showed fewer stained chromocenters in *hdl* than WT plants (Fig. 2C). These observations prompted us to assess the decompaction of chromocenters resulting from the altered heterochromatin organization. Transposable elements (TEs) such as *TA3, CACTA* and *TRANSCRIPTIONALLY SILENT INFORMATION* (*TSI*) repeats are normally repressed in the heterochromatin chromocenters region in WT plants; however, they were de-repressed in *hdl* mutants (Fig. 2D). These results indicate HDL is required for heterochromatic chromocenter maintenance.

**Figure 2.**
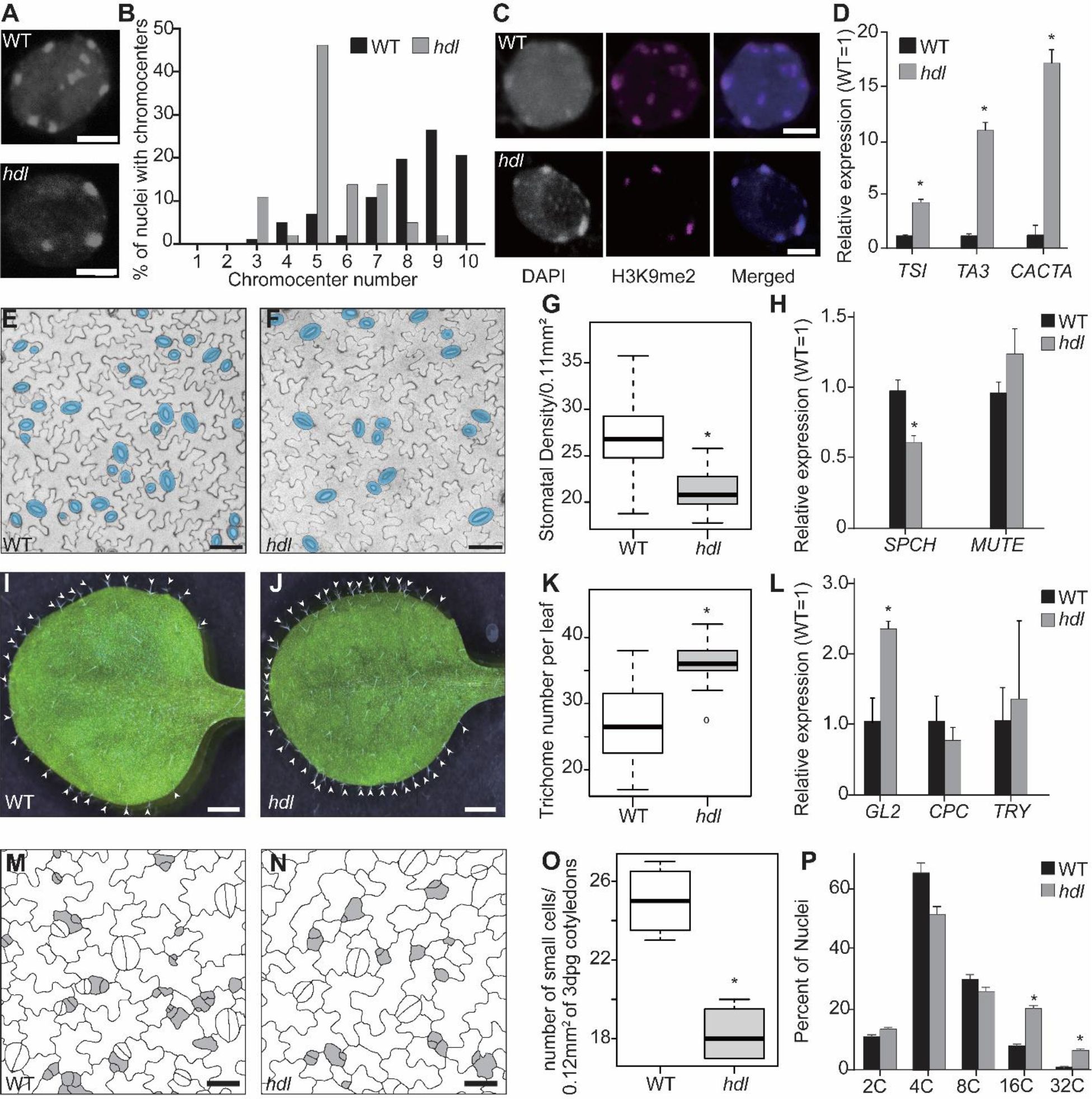
HDL is required for chromocenter organization and balances cell division and differentiation during the development. **(A-B)** Representative images and quantification of DAPI-stained chromocenters in the WT and *hdl* mutant, n=36 for each genotype). **(C)** Immunostaining of root nuclei with heterochromatin marker H3K9me2 (magenta) and DAPI (blue) in WT and *hdl* plants. **(D)** Quantitative RT-PCR (qRT-PCR) analysis of *TSI, Ta3*, and *CACTA* using 5-dpg seedlings. three biological replicates were used. **(E-G)** Stomatal density was reduced in *hdl* mutants compared to WT plants. Confocal images of 7-dpg abaxial cotyledon epidermis with cell outlines visualized using RCI2A-mCherry and stomata pseudo-colored in blue. n=20. Within each box, horizontal black lines show the median, boxes extend from the 25^th^ to the 75^th^ percentile of each group’s distribution of values; dots denote outliers. **(H)** Relative expression of stomatal regulators, *SPEECHLESS* (*SPCH*) and *MUTE*, measured by qRT-PCR. **(I-K)** Trichome number was increased in *hdl* mutants compared to WT plants. Images from the first pair of the true leaves in 12-dpg plants. n = 20. **(L)** Relative expression of trichome-related genes, GLABRA2 (GL2), CAPRICE (CPC), and TRY (TRIPTYCHON), measured by qRT-PCR. **(M-O)** Stomatal precursor cell density was reduced in the *hdl*. Precursor cells are cells smaller than 300 um^2^ (gray). Cell outline is marked with *ML1p:RCI2A-mCherry*. **(P)** *hdl* mutants had a higher portion of polyploid cells than WT. Nuclei from the first pair of true leaves were stained with propidium iodide, and ploidy was analyzed by flow cytometry. *, p<0.01 by Mann-Whitney non-parametric test. Data are medians (interquartile range). Scale bars, 2 µm in A, C, and 20 µm in E, F, I, J, M, N.

Because we had previously identified HDL as a gene highly expressed in SLGC, we also analyzed the role of HDL in leaf epidermal development. The *hdl* mutant had fewer stomata on the abaxial and adaxial leaf surfaces (Fig. 2, E to G and fig. S3M). Interestingly, trichome density in the first pair of the true leaves was increased (Fig. 2, I to K ad fig. S3N) in *hdl* compared to WT plants. Consistent with these phenotypes, *hdl* had significantly reduced expression of SPEECHLESS (SPCH), the key TF that drives cell differentiation in the stomatal lineage (Fig. 2H). However, the expression of *MUTE*, a later stomatal lineage stage gene, was not significantly different from the WT. Similarly, *hdl* had increased expression of *GLABRA* (*GL2*), which promotes trichome formation, but did not differ from WT in the expression of CAPRICE (CPC) and TRIPTYCHON (TRY) which act as negative regulators of trichome formation (Fig. 2L).

Correct leaf morphogenesis depends upon balanced cell proliferation and differentiation. Much of this balance is determined by progenitor cells in the leaf epidermis which are poised between two modes of the cell cycle, mitosis, and endoreduplication. Mitotic cells enter the diploid stomatal linage to generate more SLGCs that retain cell division potency. Cells that undergo endoreduplication differentiate into polyploid trichomes or pavement cells (*7, 17*). Therefore, we wondered whether changes in division potential and ploidy level could explain the phenotypes of *hdl* plants. Reduced stomatal density could result from reduced SLGCs division and fewer protodermal cells entering the stomatal lineage (Fig. 1A). To quantify the number of small stomatal precursor cells, we used 300μm^2^ as a cutoff and found that there were fewer small cells in 3dpg *hdl* cotyledons (Fig. 2, M and O). Also, *hdl* mutants had short roots, which could result from reduced cell division in the root meristem or reduced cell size (fig. S3, D to E and I, to J). Consistent with its reduced root length, *hdl* had a smaller root meristem and reduced number of root meristem cells (fig. S4, A and B). Both the short root and reduced root meristem size were rescued by HDLp:HDL-YFP (fig. S3, D to E and S4A). Furthermore, we used a Cyclin B1 reporter line with β-glucuronidase (GUS) and a cyclin destruction box (*CYCB1p:GUS*) to estimate cell division activity (*18*). Compared to the WT, *CYCB1p:GUS* formed fewer foci in *hdl* roots (fig. S4C) and emerging true leaves (fig. S4D), indicating that mitotic activity was reduced by loss of HDL.

Conversely, since trichomes are polyploid cells, the higher trichome density in the *hdl* (Fig. 2, I to K) may indicate a higher ploidy level in *hdl* mutants. Thus, we used flow cytometry to analyze the ploidy level and found that *hdl* had increased numbers of 16C and 32C cells compared to the WT (Fig. 2P). Overall, our data suggest that disrupted chromatin organization in *hdl* influenced meristem function by promoting the switch from cell proliferation to cell differentiation.

## HDL interacts with histone modifiers and affects histone methylation and acetylation

Heterochromatin is established through DNA methylation and histone modification. Loss of histone methyltransferases (HMTs) and histone deacetylases (HDAs) disrupt heterochromatin condensation (*19, 20*). H3K9me2 is one of the markers of heterochromatin in plants and its colocalization with HDL (Fig. 1E) prompted us to investigate whether HDL can interact with HMTs and HDAs to regulate heterochromatin formation. The *Arabidopsis* H3K9 methyltransferases KRYPTONITE/SUPPRESSOR OF VARIEGATION 3–9 HOMOLOG 4 (KYP/SUVH4), SUVH5, and SUVH6 are mediators of H3K9me deposition and function redundantly in TE silencing and heterochromatin maintenance (*21, 22*). On the other hand, KYP/SUVH5/6 can directly interact with the histone deacetylase HDA6 to synergistically regulate TE expression. Our previous HDA6 immunoprecipitation-mass spectrometry (IP-MS) dataset also showed that HDA6 interacts with HDL (*23*). Consistent with this data, <u>b</u>iomolecular fluorescence <u>c</u>omplementation (BiFC) assays also detected HDL interaction with HDA5, HDA6, HDA19, KYP/SUVH4, SUVH5, and SUVH6 in the nucleus. Conversely, HDA9 and the negative control KIP-related protein 1 (KRP1) did not produce any signal (Fig. 3, A and fig. S5). Co-immunoprecipitation (co-IP) assays also found HDL interaction with HDA6 and KYP (Fig. 3B). These data suggest that HDL functions with HDAs and HMTs to assemble and maintain heterochromatin.

**Fig.3.**
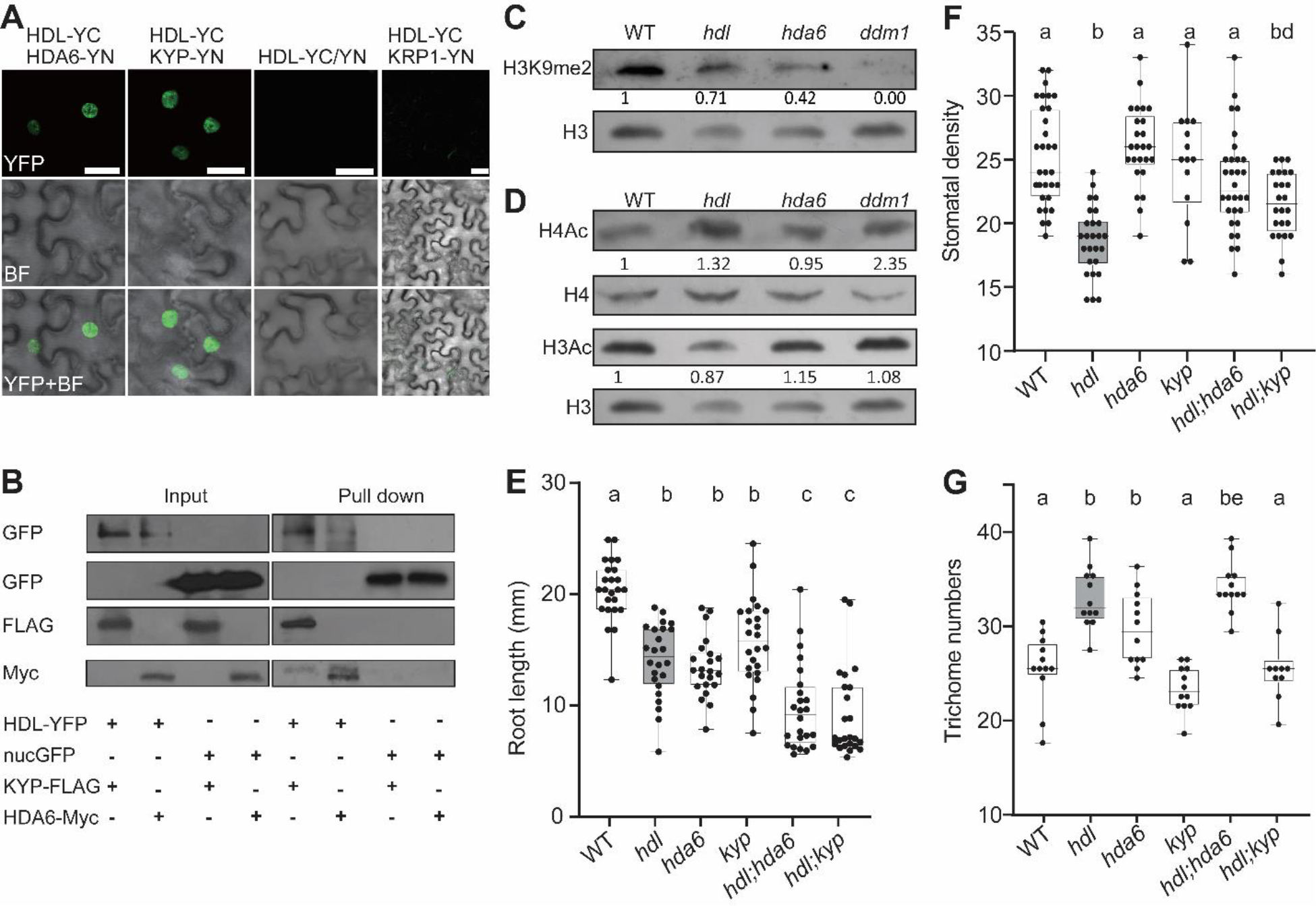
HDL interacts with histone deacetylase 6 (HDA6) and methyltransferase (KYP). **(A)** Confocal images of BiFC assays in *N. benthamiana* demonstrate the interaction of HDL-cYFP with HDA6-nYFP and KYP-nYFP. HDL-YC with empty vector and KRP1-YN were used as controls. **(B)** Western blots of Co-immunoprecipitation (co-IP) using anti-GFP antibodies revealed the interaction of HDL-YFP with HDA6-Myc and KYP-FLAG. **(C-D)** Western blots of histone methylation (H3K9me2) and acetylation (H4Ac and H3Ac) in 7-dpg seedlings of WT and *hdl* plants. Band intensities were normalized with total H3 or H4 and compared to those in the WT. n=3 **(E)** Root length quantification of WT, *hdl*, hda6, *kyp, hdl;hda6*, and *hdl;kyp*. n =22 for *hda6* and *24* for WT, *hdl, kyp, hdl;hda6, hdl;kyp*. **(F-G)** Stomata and trichome phenotypes of WT, *hdl*, hda6, *kyp, hdl;hda6*, and *hd;kyp*. More than ten leaf epidermis were examined. Letters show a significant difference by Tukey’s HSD and one-way ANOVA.

In *Arabidopsis*, SUVH4/5/6 catalyzes H3K9 methylation while HDAs remove acetylation, which prompted us to investigate whether HDL impacts histone methylation or acetylation levels. Immunoblot analysis found that *hdl* had reduced H3K9me2 level but increased level of H4Ac (Fig. 3, C and D). In contrast, there was only a marginal effect on H3Ac in *hdl*. Taken together these results demonstrate that HDL associates with H3K9me2 and, in conjunction with histone modifiers like HDA6 and KYP, participates in the establishment and maintenance of heterochromatin.

Furthermore, we also tested the genetic interaction of HDL and HDA6, KYP. Single mutants *hdl* and *hda6* had shorter roots compared to WT plants while *kyp* grows relatively normal (Fig. 3, E and fig. S6), probably because of functional redundancy with other histone modifiers such as SUVH5 and SUVH6 (*23, 24*). Double mutants of *hdl;hda6*, and *hdl;kyp* exacerbated the short root phenotype, but moderate changes in stomatal and trichome numbers, suggesting HDA6 and KYP may function with HDL differently in specific tissues or cell types (Fig. 3, F and G).

## HDL binds to both heterochromatin and euchromatin regions

To investigate the genome-wide function and *in-vivo* genome association of HDL, we performed chromatin-immunoprecipitation followed by sequencing (ChIP-seq) in the *HDLp:HDL-YFP* transgenic line. The HDL ChIP-seq data identified 6397 binding peaks, mostly in promoters (60.36%), with fewer peaks identified in introns (9.33%) or exons (8.22%) (Fig. 4A). Further analysis of the HDL binding targets showed enrichment in protein-coding (88.4%) and transposable element genes (6.19%). Within the top 1% of HDL binding peaks, 40 of them were located in transposable element genes (63.49%) while 20 of them resided in protein-coding genes (Fig. 4B).

**Figure 4.**
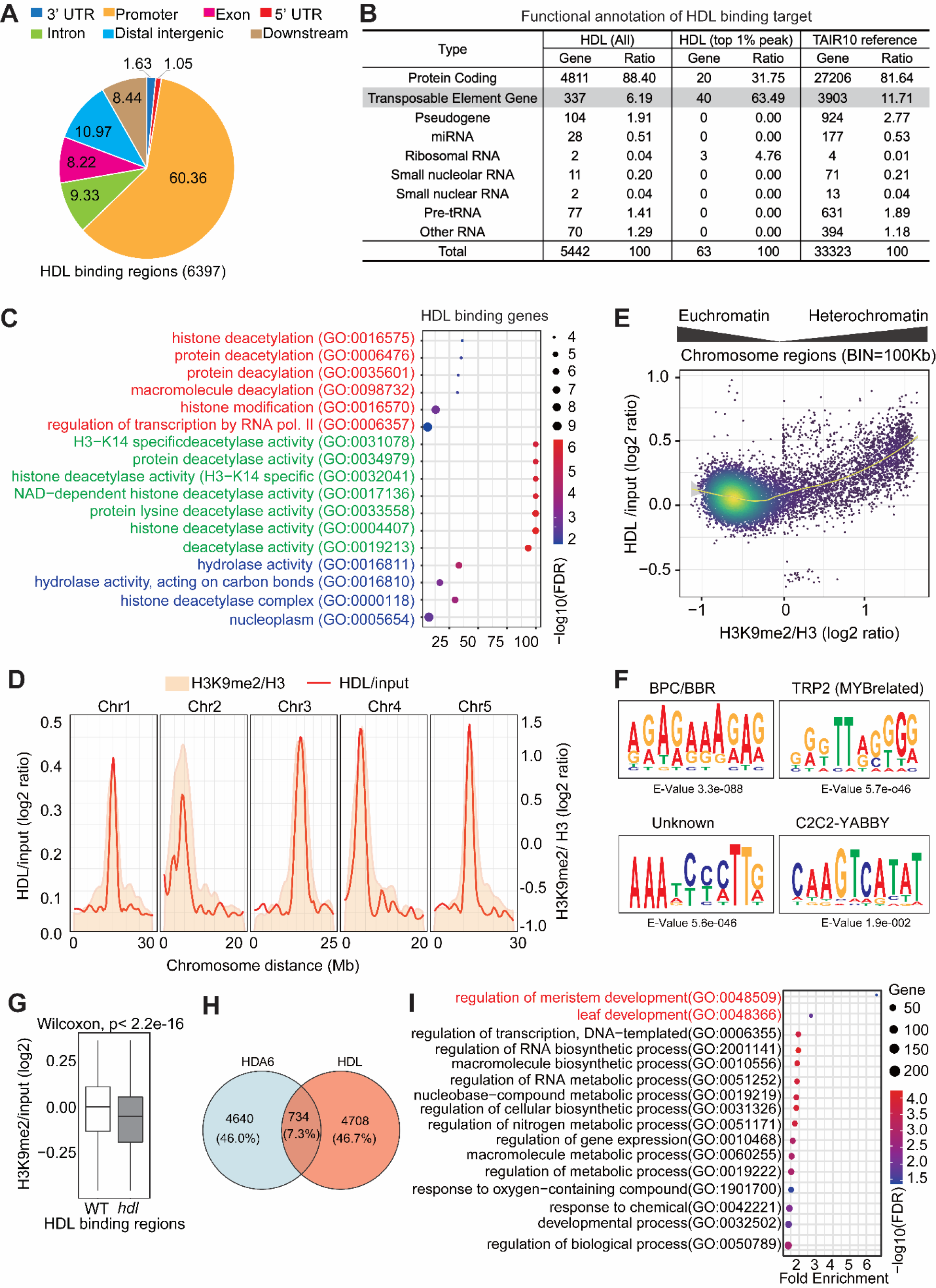
HDL is present in both heterochromatin and euchromatin regions with development implications. **(A)** ChIP-seq analysis reveals a significant enrichment of HDL binding in promoter regions. Promoter regions refer to the 3 Kb upstream of the transcription start site, while downstream regions cover the 1 Kb segment extending beyond the gene’s end. **(B-C)** Functional annotation of HDL ChIP-seq peaks reveals that the top 1% of binding peaks are predominantly located within transposable element gene regions and the enriched protein-coding genes were associated with histone deacetylation and modification in the GO enrichment analysis. **(D)** The enrichment of HDL corresponded to the genome-wide distribution of H3K9me2. The red line represents the HDL/input ratio, and the yellow area indicates the H3K9me2/H3 levels across five chromosomes. **(E)** Genome-wide average HDL/input ChIP-seq levels are plotted against H3K9me2/H3 ChIP-seq levels. HDL preferentially associated with regions with higher H3K9me2/H3 levels, while numerous sites of HDL association were also present in euchromatin regions with lower H3K9me2/H3 levels. **(F)** Top motif categories identified from HDL-ChIP analysis. An analysis of 100bp sequences centered around the HDL binding summit using MEME (*1*) revealed that the most prominent motif corresponded to GAGA-binding factors. **(G)** The H3K9me2 levels in HDL binding regions awee reduced in *hdl* mutants compared to WT plants. **(H)** A Venn diagram illustrates the overlap of co-regulated genes of HDL and HDA6 (*2*). **(I)** The co-binding genes of HDL and HDA6 exhibited significant enrichment in GO terms associated with meristem and leaf development.

The top 1% HDL binding peaks were highly enriched for Gene Ontology (GO) terms related to histone deacetylation, histone modification, and regulation of transcription (Fig. 4C). Consistent with immunostaining data (Fig. 1E), our ChIP-seq analysis revealed that HDL was preferentially enriched in regions with high levels of H3K9me2 and in centromere (Fig. 4, D and fig. S7). Although HDL showed a high preference for heterochromatin association, it did also associate with euchromatin as revealed by the dot density plot where we also observed HDL enrichment is in regions with H3K9me2/H3 level lower than 0 (Fig. 4E). This suggested a possible role of HDL in transcriptional regulation. Functional annotation of the HDL-bound promoter targets found enrichment of genes associated with trichome patterning and flavone biosynthesis-related processes, indicating a role of HDL in leaf development (fig. S8, A and B and Data S2).

To identify the specific sites of HDL chromatin association, 100 bp sequences at the peaks of HDL chromatin association were screened using motif analysis of large nucleotide datasets (MEME-ChIP, E-value <0.05). This analysis identified a significantly enriched motif related to BARLEY B RECOMBINANT/BASIC PENTACYSTEINE (BBR/BPC), also known as the GAGA motif, comprised of tandem repeats of GA (Fig. 4F). The other most enriched motifs were TRP2 (MYB related) and C2C2, which also contains GA repeats. C2C2 motifs are bound by the TFs containing zinc finger or helix loop helix domains such as High Mobility Group (HMG) proteins, which are involved in carpel cell fate (*25*). Recent studies have demonstrated that HDA6 and KYP-occupied genomic regions are enriched in GA repeats. (*24, 26*), supporting the HDL interaction with HDA6 and KYP. The enrichment of the BBR/BPC motif particularly drew our attention because the BBR/BPC motif is bound by the plant-specific BBR/BPC family of TFs that regulate *Arabidopsis* organogenesis by changing the transcriptional regulatory network of homeobox genes (*27, 28*). Recently, BBR/BPC motifs have been found to bound by SPCH and MUTE, the two master bHLH regulators of the stomatal lineage, and BPC acts as a trans-acting factor to suppress entry into proliferative divisions of the stomatal lineage (*29*). Hence, this suggests that enrichment of BBR/BPC motifs may direct HDL to the same genomic regions as HDAs and KYP to change the methylation level at HDL-associated sites (Fig. 4G), thus leading to chromatin modifications that determine the specificity of cell fate.

## HDL suppresses *HDA6* and *KYP*

Based on the interaction of HDL with HDA6 (Fig. 3), we hypothesize that they may co-regulate transcription by associating with the same regions and modifying chromatin architecture. By comparing the HDL binding regions with previously published ChIP-seq data of HDA6 (*26*), we found that 734 (7.3%) of genes shared the binding of HDL and HDA6 (Fig. 4H, and Data S3).

The GO-term analysis of these co-binding genomic sites indicated an enrichment of genes involved in the regulation of meristem and leaf development (Fig. 4I). It has been shown that the GAGA-binding factor, BPC6, recruits polycomb-repressive complex 2 to Polycomb responsible elements and maintains genes in a repressed state (*30*). Therefore, GAGA-motif-enriched HDL may also function with other chromatin regulatory proteins to repress the transcription of developmental genes in leaf epidermis. Consistent with such a hypothesis, HDL was associated with genes related to histone deacetylation (Fig. 4, C, and Data S4) and may function in compacting chromatin. To investigate the functional correlation between HDL and *HDA6*, or *KYP*, we performed a dual luciferase reporter assay using *35S:HDL* as effector, and *HDA6p:LUC* and *KYPp:LUC* as reporters. The results showed that HDL repressed *HDA6* and *KYP* promoter activity (fig. S9). These results indicate that while HDL and HDA6/KYP function together in compacting chromatin, the HDL may form a feedback loop to regulate the expression of *HDA6* and *KYP*, thereby controlling heterochromatin state. Collectively, these results show that HDL associates with H3K9me2 and functions together with histone modifiers such as HDA6 and KYP to establish and maintain heterochromatin in leaf epidermal development.

## HDL regulates chromatin accessibility and alters the epidermal cell fate

The interaction of HDA6 and KYP with HDL (Fig. 3), and the HDL enrichment at heterochromatin (Fig. 4) led us to hypothesize that HDL may regulate chromatin accessibility. To analyze changes in chromatin accessibility we performed Assay for Transposase Accessible Chromatin (ATAC) coupled with high throughput sequencing. Our results show that the most accessible chromatin regions in *hdl* were upstream of transcription start sites (TSS) (Fig. 5, A-B), suggesting that HDL functions in transcription regulation. To further narrow down the sites of HDL action, we compared our ChIPseq and ATAC datasets and found that HDL-associated regions also had higher chromatin accessibility in *hdl* (Fig. 5, C-D). To determine whether this altered chromatin accessibility correlates with transcriptional changes, we performed transcriptomic analysis after using fluorescence-activated cell sorting to collect epidermal cells of WT and in *hdl* plants. Although the overall transcriptome does not show drastic changes between WT and *hdl* plants (fig. S10), genes related to stomatal development were significantly downregulated in *hdl* (Fig. 5E, and Data S5). Also, we found that the number of cells expressing SPCH was significantly reduced in *hdl* compared to the WT (Fig. 5F, and G), even though SPCH-bound and stomata-related genes showed higher chromatin accessibility at the promoter regions (fig. S11, A, and B, Data S6). We then used an integrated genome browser to visualize the HDL chromatin association, accessibility, H3K9me2 level, and transcription of genes related to stomata, trichomes, and epigenetic regulators. Our results showed that the H3K9me2 level of those genes was reduced which leads to an increase in the accessibility in *hdl* mutants compared to WT (fig. S12).

**Figure 5.**
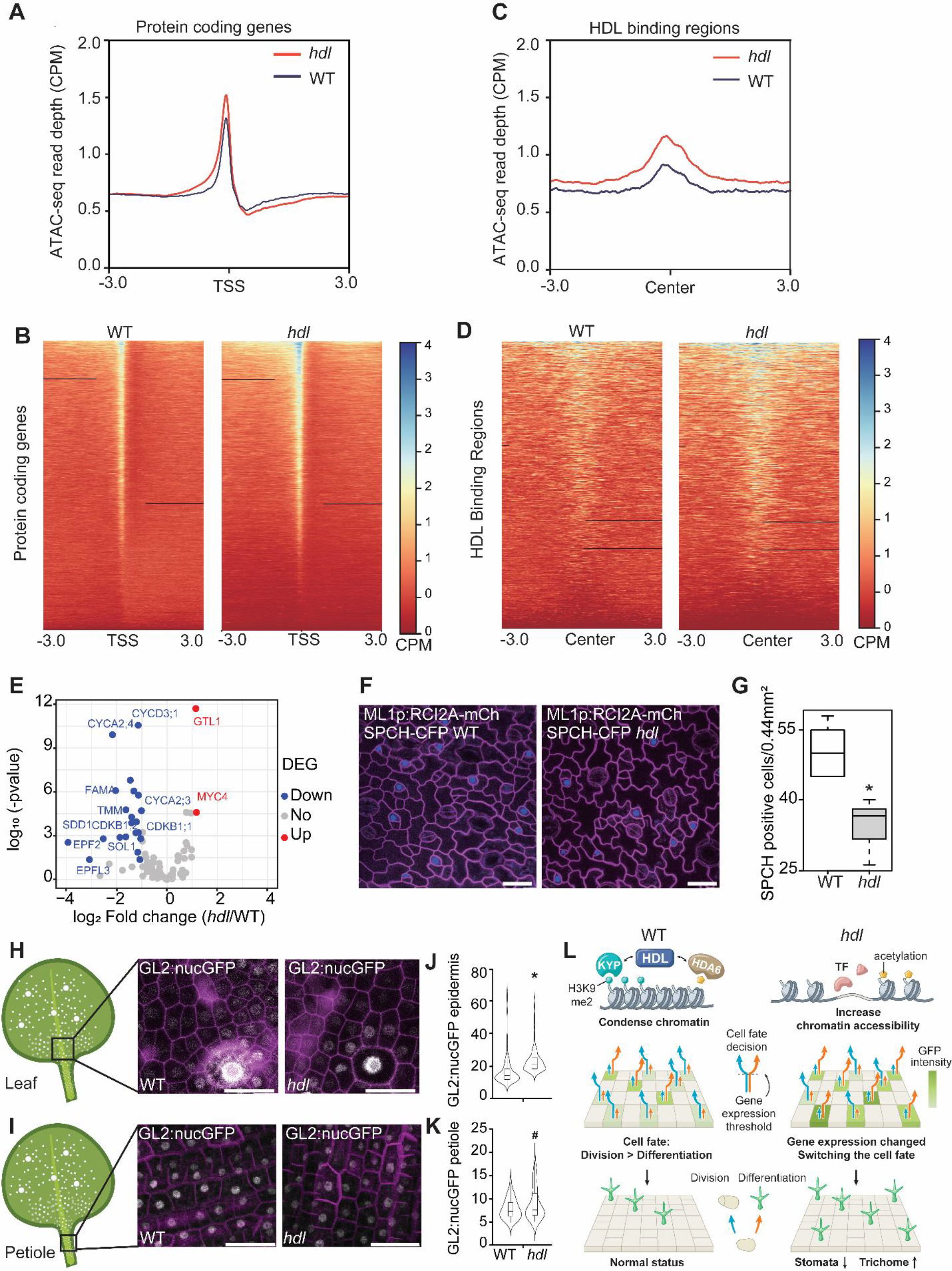
HDL regulates chromatin accessibility and manipulates epidermal cell fate decisions. **(A)** In *hdl* plants, chromatin accessibility is higher in the regions before Transcriptional start sites (TSS) compared to WT plants. ATAC-seq read depths from WT and *hdl* were plotted against all protein-coding genes based on the TAIR10 annotation. **(B)** Heatmaps of ATAC-seq data for all protein-coding genes displayed in (A). Each row represents an individual protein-coding gene. **(C)** The loss of HDL leads to an increase in chromatin accessibility within HDL-binding regions. ATAC-seq read depths from WT and *hdl* were plotted against HDL binding regions, with the binding sites as the reference point for region alignment (in kilobase pairs). **(D)** Heatmaps of ATAC-seq data for all HDL binding regions shown in (C). Each row corresponds to a distinct HDL-binding region. **(E)** Differential expression analysis of stomatal-related genes found down-regulation in *hdl* compared to WT plants. Red dots represent up-regulated genes (log2Fold Change (*hdl*/WT) >2, p-value < than 0.01), while blue dots represent down-regulated genes. **(F-G)** Confocal images and quantification of SPCH-CFP expression in the abaxial epidermis showed a reduced number of cells expressing SPCH-CFP in *hdl* compared to WT plants. *, p<0.01 by Mann Whitney non-parametric test. **(H-I)** Confocal images were captured from the adaxial true leaf epidermis and petiole of both WT and *hdl* plants expressing *GL2p:nucGFP*. For the epidermis, n=291 for WT and n=266 for *hdl*, while for the petiole, n=35 for WT and n=56 for *hdl*. **(J-K)** Quantification of *GL2p:nucGFP* signal in (H) found higher *GL2* levels in *hdl* mutants than in WT plants at the base of the true leaves. Additionally, the variance of nucGFP levels was significantly greater in *hdl* mutants compared to WT at the petioles (H-I). Mean comparison: *, p<0.01 by Mann-Whitney non-parametric test. Variance comparison: ^#^, p<0.01 by Bartlett’s test. Scale bars, 40 µm in G, I, and K. **(L)** In WT plants, HDL interacts with HDA6 and KYP to condense chromatin, thereby regulating gene expression and maintaining the balance between cell division and differentiation. The loss of HDL results in altered chromatin accessibility, impacting transcriptional activity in the leaf epidermis and increasing variability in GL2 expression, as indicated by GFP intensity. This ultimately leads to changes in leaf epidermal patterning.

During the early stages of development, stem cells may undergo transcriptional bursts associated with specific fates, and the frequency of such bursts may cause expression variability (noise) in gene expression. It has been proposed that chromatin structure influences gene expression noise (*31, 32*). Thus, we measured the gene expression variability of *GL2* as it is closely related to the *hdl* trichome phenotype. We quantified *GL2* reporter expression in early trichome precursor cells or the petiole cells which undergo rapid cell division from the adaxial epidermis of the emerging true leaf and found that *GL2* intensity was significantly higher in *hdl* than WT (Fig. 5H, and J). However, the variation of GFP intensity is larger in *hdl* than WT in the petiole region (Fig. 5I, and K), indicating the increased variability of cell fate choice. Furthermore, *ddm1* mutants with disrupted chromocenters exhibited an excess of trichomes and reduced stomatal density, phenotypes similar to that of *hdl* mutants (fig. S13).

Here, we reveal the role of HDL, a previously uncharacterized homeodomain-containing protein, in the organization of chromatin. As leaf epidermal development begins, protodermal cells with well-organized chromatin tend to rapidly undergo cell division to produce more stomatal precursor cells. When HDL is absent, looser chromatin organization increases the accessibility of many genes. This leads to downregulation of genes related to stomatal development and more variable expression of *GL2*, a gene crucial for trichome formation. Consequently, this chromatin-mediated influence enhances the chance of epidermal precursors adopting a trichome cell fate, leading to a critical shift from cell proliferation to differentiation (Fig. 5L). Our findings suggest a significant role of chromatin organization in shaping the transcriptional activity of leaf epidermal precursors. In addition to its impact on transcription regulation, the alternation of the cell cycle also has impacts on cell fate determination. For example, the cell cycle inhibitor SIAMESE-RELATED1 (SMR1) plays a role in terminating the self-renewal potential of SLGCs, a process that also requires CYCLIN A (CYCA) and CYCLIN-DEPENDENT KINASE B1 (CDKB1) (*33*). Additionally, SMR4 orchestrates the differentiation of guard mother cells by decelerating cell cycle progression (*34*). We also observed the downregulated expression of *CYCA*s, *CDKB*s, and *CYCD3;1* in *hdl* (Fig. 5E), which underlines the importance of the cell cycle in the determination of cell fate. Nevertheless, the intricate interplay among chromatin structure, cell cycle, and cell fate will require further investigation to be fully understood.

## Acknowledgments

We thank Dr. Masaaki Umeda at Nara Institute of Science and Technology for his kind input at the beginning of this project. We thank Dr. S.-Y. Chen and Y.-T Lu at the Institute of Biomedical Sciences, Academia Sinica for assistance with ATACseq experiments and T.-W. Tai and C.-C Tai at flow cytometry core facility for assisting with cell sorting. Dr. D. Bergmann (Stanford University and Howard Hughes Medical Institute) for providing stomata-related constructs, and the Arabidopsis Biological Resource Center for providing seeds. We would like to extend our gratitude to the core facilities at the Institute of Plant and Microbial Biology, Academia Sinica. Special thanks to Dr. Y.-F. Tseng at flow cytometry core facility for assisting ploidy analysis; S.-J. Chou and A.-P. Chen for assisting with the Illumina library preparation and sequencing; Dr. W.-D. Lin and Y.-I. Lin for assisting with sequencing data analysis; J.-Y. Huang at the Live-Cell Imaging Division, Cell Biology Core Lab for advice on using the Leica STELLARIS 8 and Zeiss LSM880 confocal microscopes; M.-J. Fang and M.-L. Cheng at the Genomic Technology Core Lab for the DNA sequencing service. We thank Dr. Paul Verslues (IPMB, Academia Sinica) for his suggestions on this manuscript.

## Author contributions

A.A., C.K., and C.K.H. conceived the study and designed the experiments. C.K. and T.C. conducted the ATAC-seq experiment and analysis. F.H. and Y.F. carried out the HDL-ChIP assay and BiFC assay. S.Y. performed the H3 and H3K9me2-ChIP experiments. H.L. executed the pull-down assay. A.A. conducted all other experiments and analyses. C.K. conducted the integrated data analysis with all the sequencing datasets. K.W. supervised F.H. and Y.F. A.A. and C.K.H. wrote the manuscript with feedback from C.K., F.H., and K.W.

## Data and materials

All data are available in the main text or the supplementary materials. All materials (seed stocks, plasmids) are available upon reasonable request.

