## Supplementary Material for "The HOMEODOMAIN-Like protein HDL mediates chromatin organization and rewires leaf epidermal patterning"

**The PDF file includes**

Materials and Methods

Figs. S1 to S13

Table S1

References

**Other Supplementary Material for this manuscript includes the following:**

MDAR Reproducibility Checklist

Data S1 to S6

### Materials and methods

#### Plant material and growth condition

*Arabidopsis thaliana* Col-0 was the WT used in all experiments, and all transgenic lines were created in this background. The *hdl* mutant, SALK-052138, was obtained from ABRC. The *hda6* (*axe1-5*), *kyp* (*1*) and *ddm1* mutants have been previously described (*2, 3*). The plant reporter lines used in this study include AtML1pro:RCI2A-mCherry, SPCHp:SPCH-CFP (*4*) CENHp:CENH3-GFP (*5*). The primers used to generate the constructs are listed in Table S1. Ethanol (EtOH) sterilized seeds were sown on ½ Murashige and Skoog (MS) medium plates and stored at 4°C for 2 days of stratification. After that plates were transferred to a plant tissue culture room at 22°C under a 16-hour light/8-hour dark cycle (light intensity of 80-100 µmole $m^{-2}$ $s^{-1}$, wavelength 400-720 nm, bulb type: florescencent tube). Plant materials used for phenotyping and the extraction of mRNAs and proteins were grown on 1/2 MS plates for seven days, unless until specified.

#### Plasmid constructions

For reporter and luciferase assay, constructs were created by Gateway (Invitrogen) cloning with pENTR 5'-TOPO, pENTR-D-TOPO and pDONR P4-P1R plasmids as backbone. To facilitate the cloning process, we modified the EcoR1 site (NEB) in pENTR 5'-TOPO to NotI and AScI sites; the resulting plasmid was named as pENTR5'S. Genes were cloned into pENTR-D-TOPO plasmids by using conventional TOPO isomerase (Invitrogen) or by restriction enzyme (RE) digestion with NotI and AScI (NEB) depending on the primer design list Table S1. **HDLp:HDL-YFP** was generated using pENTR 5'-TOPO-HDLp, pENTR/D-TOPO-HDL and R4pGWB540 (*6*) (Nakagawa et al., 2008). **35S:HDL-YFP** was obtained using pENTR/D-TOPO-HDL and pH35GY (Kubo et al., 2005). **H2AW.6p:H2AW.6-CFP** was obtained by using pENTR 5'S-H2AW.6p, and pENTR/D-TOPO-H2AW.6 and R4pGWB443. **GL2p:nucGFP** was cloned by using pENTR/D-TOPO GL2p with pGGN. To generate ***hdl-crispr*** lines, the two guide RNA (gRNA) sequences (Table S1) were introduced into plasmids containing AtU6-26-sgRNA by BsaI (NEB). These constructs were then introduced into pCAMBIA1300-221 containing YAOp:Cas9 by SpeI and NheI (*7*).

For subcellular localization, co-localization, and complementation assays, the constructs were introduced into the *Arabidopsis* via *Agrobacterium tumefaciens* (GV3101) via the floral dip method (*8*).

**The phenotype of roots, stomata, and chromocenters**

**For the root growth**, the plants were grown in ½ MS medium under long-day conditions (16-hour light and 8-hour dark) in the vertical direction for four days and then transferred to new plates for 3-5 days to track and measure the root length. **For the stomatal phenotype**, plants were grown horizontally and seedlings were fixed overnight in a 7:1 solution of ethanol:acetic acid. The samples were washed with Milli-Q water and subjected to softening in 1M KOH until they became entirely transparent. Subsequently, they were mounted in Milli-Q water for observation using a Leica DM2500 LED microscope equipped with a DIC prism (Leica Microsystem). **For measuring small cell density**, 3-dpg old plants abaxial cotyledons were used. **For the quantification of chromocenter number and size**, cotyledons from 3-day-old plants were fixed in a solution of 7:1 (ethanol: acetic acid), followed by two washes with dd$H_2O$. Subsequently, they were stained with DAPI for 5 minutes and observed directly under the Leica Stellaris Stellaris 8 (Leica Microsystem) microscope. The StarDist plugin in Fiji ImageJ was employed to detect and quantify the nuclei.

**Microscopy**

Protein subcellular localization and plasma member marker were monitored using a Leica STELLARIS 8 or a Zeiss LSM880 (Carl Zeiss system). For trichome analysis, the first pair of true leaves were imaged with Stereo Fluorescent Microscope (Zeiss Lumar V12). The *GUS* staining signal was monitored with a Upright Fluorescent Microscope (Zeiss Axio Imager Z1). The fluorescent proteins, CFP, GFP, YFP, and mCherry were excited with 458-nm, 488-nm, 514-nm and 561-nm lasers, respectively.

**GUS staining**

WT or *hdl* plants carrying the *CYCB1p:GUS* reporter construct (*9*) were grown vertically on solid ½ MS medium for 7 days. Subsequently, the seedlings were immersed in ice-cold acetone (90%) for 10 minutes, followed by a rinse with dH2O. Next, they were submerged in GUS staining solution (100 μM K3Fe(CN)6, 100 μM K4Fe(CN)6, 50 mM phosphate buffer, and 1 mM X-GLUC in dimethyl formamide). Vacuum infiltration was performed twice for 10 minutes each. The seedlings were then incubated at 37°C for 3-5 hours. After incubation, the samples were placed at 4°C overnight in 80% ethanol. On the following day, the samples were washed with methanol and acetic acid solution (7:1) for 15 minutes at room temperature and subsequently mounted on slides in ethanol for imaging using an Imager Z1 microscope (Zeiss).

**Real-Time qPCR analysis**

For the detection of mRNA expression, total RNA was extracted from plant tissues or root tips (up to 5 mm in length) for quantifying mRNA expression, using the RNA Plus Mini Kit (LabPrep). Complementary DNA (cDNA) was synthesized from 1-5 µg of total RNA using SuperScript II transcriptase (Thermo Fisher Scientific). The qRT-PCR reaction mixture comprised a cDNA template (100 ng), specific primers, and SYBR Green Master Mix (Applied Biosystems). Quantitative real-time PCR was conducted using the QuantStudio 12K Flex Real-Time PCR cycler system (Applied Biosystems). The primers used for qRT-PCR analysis can be found in Supplemental Table S2.

**Flow cytometry**

To measure the ploidy levels of *hdl* and WT plants, the first pair of true leaves from 12-day-old seedlings were finely chopped using a sharp blade in a 55-mm plastic petri dish containing 250 µl nuclei extraction buffer (200 mM Tris, 4 mM $MgCl_2$-$6H_2O$ and 0.5% Triton X-100) (Sysmex, Cystain PI Absolute P) for 1 minute. Nuclei were then stained with 1000 µl staining buffer (50 µg $ml^{-1}$ propidium iodide and RNase A, with excitation at 561nm). The mixture was then passed through a Partec 50-$\mu m$ Cell Trics disposable filter and incubated on ice for 30 minutes in the dark. The samples were subsequently analyzed with a flow cytometer (Beckman Coulter CytoFLEX Cell Analyzer).

**Luciferase reporter assay**

For the transcriptional activity assay, the reporters (HDA5p:LUC, HDA6p:LUC, HDA9p:LUC, HDA15p:LUC, HDA17p:LUC, KYPp:LUC, and ABO3p:LUC), and the effectors (HDLp:HDL-YFP, or 35Sp:nucGFP) were transilently expressed in tobacco leaves. The activities of co-expressed Renilla LUC is used to normalized the activities of LUC (luciferase) reporter. The experiment was conducted independently three times.

**Slide preparation and Immunostaining**

Slide preparation and sample fixation were performed as described (*10, 11*). To block non-specific binding, the slides were incubated with 5% BSA for 30 minutes at 37°C. Next, primary antibodies (1:200 dilution; anti-GFP Abcam, ab290; anti-H3K9me2, Abcam, ab1290) were applied and left overnight at 4°C. The slides were subsequently washed three times with 1X PBS and incubated with secondary antibodies (1:100 dilution; Goat Anti-Mouse IgG H&L (Alexa Fluor 488), Abcam, ab150113; Goat Anti-Rabbit IgG H&L (Alexa Fluor 555), ab150078) for 1 hour at 37°C or 3 hours at RT. After additional washing with 1X PBS, the slides were counterstained with DAPI and examined using a Leica STELLARIS 8 microscope.

For quantifying chromocenters, seedlings were fixed in 4% paraformaldehyde for 1 hour and rinsed with ddH2O. Root nuclei were then isolated as described above, mounted with DAPI, and observed using a Leica STELLARIS 8 microscope.

**RNA-seq analysis**

7 dpg seedings of reporter lines were used for protoplast isolation and FACS as described in (*12*). FACS was performed on Aria IIIu (BD biosciences) with an 100 mm nozzle. Approximately 20,000 mCherry positive cells/sample were sorted into 350 µl RNA extraction buffer and total RNA were extracted using RNeasy Micro Kit (QIAGEN). RNA from three biological replicates was sequenced separately, and the sequence libraries were built using the Illumina TruSeq RNA library preparation protocol. The libraries were sequenced on the NovoSeq PE150 platform using a paired-end scheme (2 x 150 bp). Reads were mapped to the TAIR10 of the *A. thaliana* genome using the Qiagen CLC genomic workbench (https://digitalinsights.qiagen.com/) with default settings. Reads that were mapped to multiple regions were discarded. Calculations to identify differentially expressed genes (DEGs) were also performed by DESeq2. Genes with at least a 2-fold- change in expression ($P<0.05$) were considered to be differentially expressed.

**Chromatin Immunoprecipitation assay (ChIP)**

The ChIP assay was conducted as previously described (*13, 14*). In brief, chromatin extracts were prepared from seedlings that had been treated with 1% formaldehyde. Subsequently, chromatin was sheared to an average length of 500 bp using sonication, and protein-DNA complexes were immunoprecipitated using specific antibodies against YFP (Abcam, catalog no. ab290) or H3K9me2 (diagenode, C15410060). The DNA that had been cross-linked to the immunoprecipitated proteins was subsequently reversed, followed by analysis using RT-PCR.

### ChIP-seq and data analyses

To ensure an adequate starting DNA amount for library construction, 2 ng of DNA from the ChIP was utilized. The ChIP DNA was initially subjected to qPCR testing and then employed for ChIP-seq library preparation using NEBNext® Ultra™ II DNA Library Prep kit (cat no. E7645), according to the manufacturer's recommended protocol. The Novoseq PE150 was used for high throughput sequencing of the ChIP-seq libraries. Raw sequence data was processed using the GAPipeline Illumina sequence data analysis pipeline. The reads were then mapped using Bowtie2 with reference to the *Arabidopsis* genome (TAIR10) (*15*). The HDLp:HDL-YFP/*hdl* line was used for ChIP-seq experiment. Over 30 million paired reads were used for two-line analysis (pair-end 150 bp). The alignments were first converted to Wiggle (WIG) files using deepTools, then imported into Integrated Genome Viewer (IGV) (*16*) for visualization. The MACS2 (*17*) was used to identify ChIP-enriched peaks, and the distribution of the ChIP binding peaks was analyzed with ChIPseeker (*18*) high-read random *Arabidopsis* genomic region subset (1,350,000 regions) was used to represent the ratio of the total *Arabidopsis* genomic regions. To identify enriched motif sites, a 100 bp sequence encompassing each peak summit (50 bp upstream, 50 bp downstream) was obtained and searched for DNA motifs using MEME-ChIP with the default parameters (*19*). The HDL:YFP ChIP-seq short read data have been submitted to the NCBI Gene Expression Omnibus (GEO) database (GSE223849)

### Phylogenetic analysis

HDL protein sequences were retrieved from the TAIR website (https://www.arabidopsis.org/). The HDL protein sequences were blasted against individual species of the algal genome to find homologs in green algae (https://phycocosm.jgi.doe.gov/phycocosm/home). The HDL homologs from other plant species were obtained from Phytozome v13 using the HDL protein sequence as a query (https://phytozome-next.jgi.doe.gov/). Predicted homologs were again blasted to verify the conserved domain in the protein sequences. All the sequences were then submitted to MAFFT or MUSCLE (https://ngphylogeny.fr/tools/tool/268/form) for multiple sequence alignment (MSA). The tree was built using the IQ tree web server (http://iqtree.cibiv.univie.ac.at) with the maximum likelihood method. The model was set as JTT and 100 bootstrap replications. The tree was edited in FigTree.v1.4.4 software.

**Imaging processing**

Root meristem images were analyzed using ImageJ (https://imagej.nih.gov/ij/download.html). Images were taken with a Zeiss LSM880 microscope (40X H2O). Meristem cell numbers were quantified from the Quiescent Center (QC) to the first cell of the differentiation zone (double the size of the cortex cell compared to the previous one).

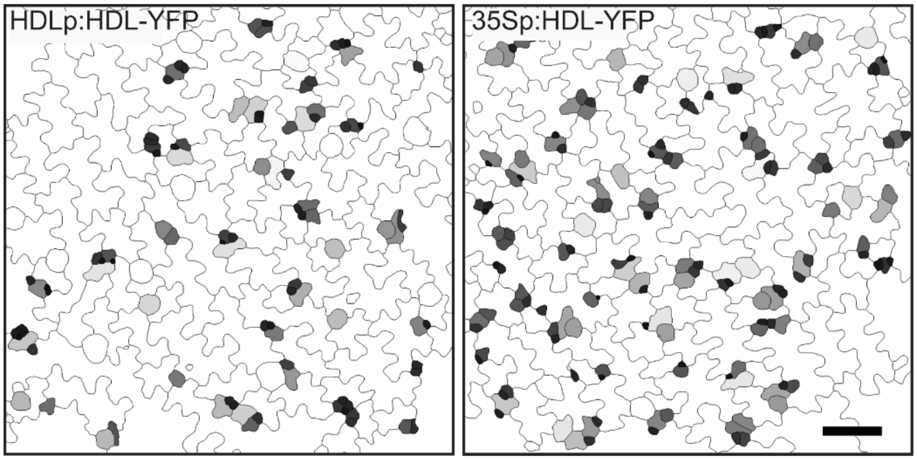


**Figure S1. Overexpression of HDL results in an increased number of small cells.** Cell size-based segmentation of transgenic lines of HDL with native or constitutive overexpression (35S) promoter. Cells labeled in gray and light gray are smaller than 300 µm$^2$: scale bar, 50 µm.

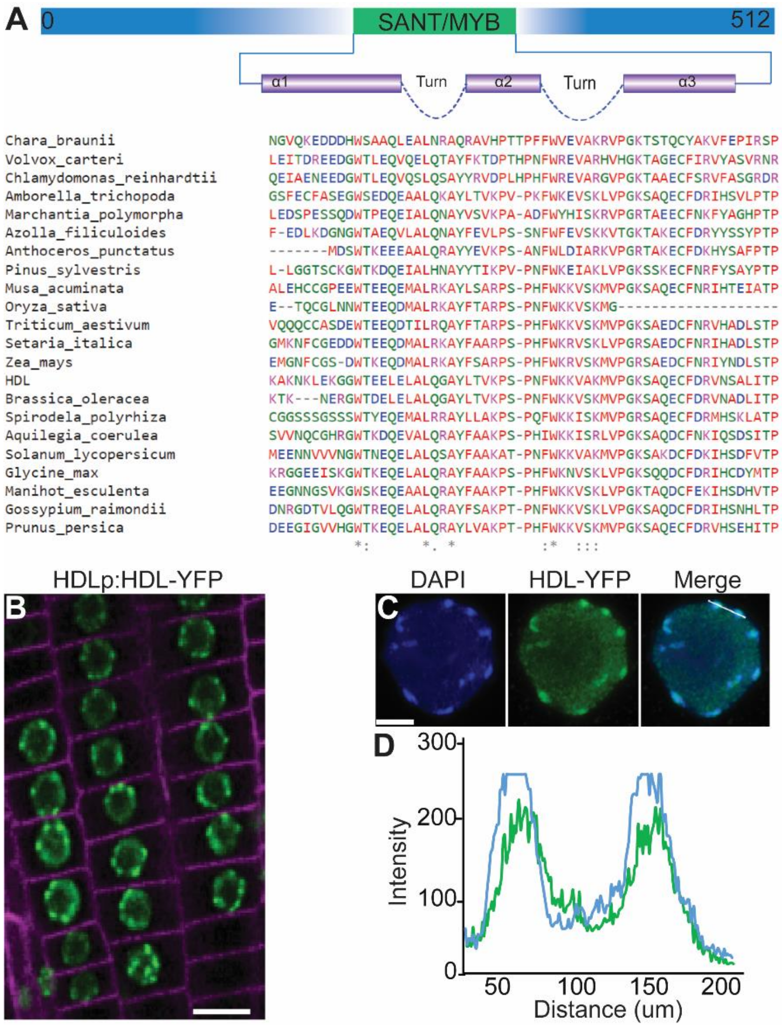


**Figure S2. HDL contains a putative DNA binding domain and is associated with heterochromatin. (A)** The domain analysis of HDL protein indicated the presence of a SANT/MYB-like DNA-binding domain from 255 aa to 307 aa. SANT/MYB consists of three alpha-helices formed in a helix-turn-helix motif. The HDL protein domain was predicted by the simple molecular architecture research tool (SMART, http://smart.embl-heidelberg.de/). Multiple sequence alignment of HDL SANT/MYB domain regions with its homologs in other plant species. The asterisks indicated conserved amino acids. **(B)** Confocal images of HDLp:HDL-YFP (green) in 5-dpg root meristem. The plasma membrane is visualized with ML1p:RCI2A-mCherry (magenta). **(C-D)** HDL colocalized with DAPI-stained chromocenters. Fixed root nuclei were immunostained with DAPI for DNA and GFP antibody for HDL-YFP. Scale bar, 10μm in B, and 2 μm in C.

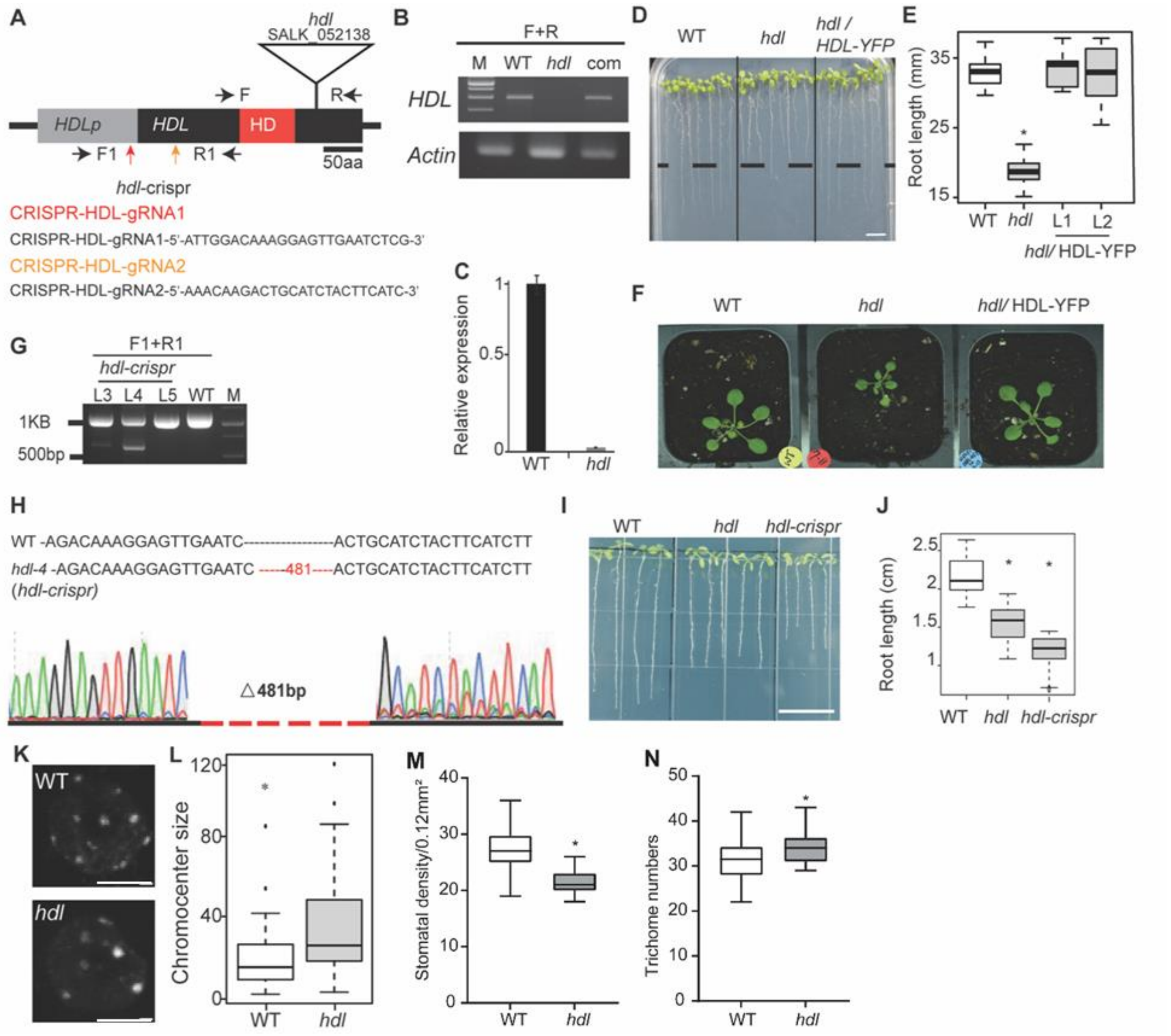


**Figure S3. Isolation and phenotype of *hdl* mutants. (A)** Schematic representation of T-DNA and CRISPR guide RNA (gRNA) insertion sites in HDL gene. The gray box denotes the promoter, the black boxes represent the CDS (Coding Sequence) region of HDL, and the red box signifies the homeodomain. The T-DNA insertion site is marked with a triangle, and CRISPR gRNA target sites are indicated by orange and red arrows. **(B)** Analysis of HDL RNA transcripts in WT, *hdl* mutants, and complementation line by RT-PCR. 7-dpg whole seedlings were used to detect HDL transcript. **(C)** The relative expression levels of HDL in WT and *hdl was* detected by qRT-PCR using 7-dpg seedlings with the WT expression level set as 1 for reference. **(D)** 2-week-old plant phenotype of WT, *hdl*, and complementation lines, *hdl*/HDL-YFP. **(E)** Root length quantification from (D). n =21 roots for all genotypes. **(F)** Phenotypic comparsion of WT, *hdl*, and complementation line at 21-dpg in soil. **(G)** Analysis of T1 generation in *hdl-crispr* lines using primers set F1 and R1. The expected deletion size is 481 bp, as shown in (H). **(H)** Sequence alignment and base pair (bp) deletion of *hdl-crispr* (line 4). The number indicates the total bp deletion. **(I-J)** Root length phenotype and quantification in 8-dpg WT, *hdl,* and *hdl-crispr* plants. n =21. **(K)** Representative confocal images of DAPI-stained chromocenters in WT and *hdl* nuclei. **(L)** Quantification of chromocenters size from (K). n =37 and 32 nuclei. **(M-N)** Stomata, and trichome density quantification from same adaxial true leaf, n= 20 and 30 true leaves respectively. $p<0.01$ by Mann Whitney non-parametric test. Scale bar, 5 mm in (D), 20 mm in (I), 5µm in (K).

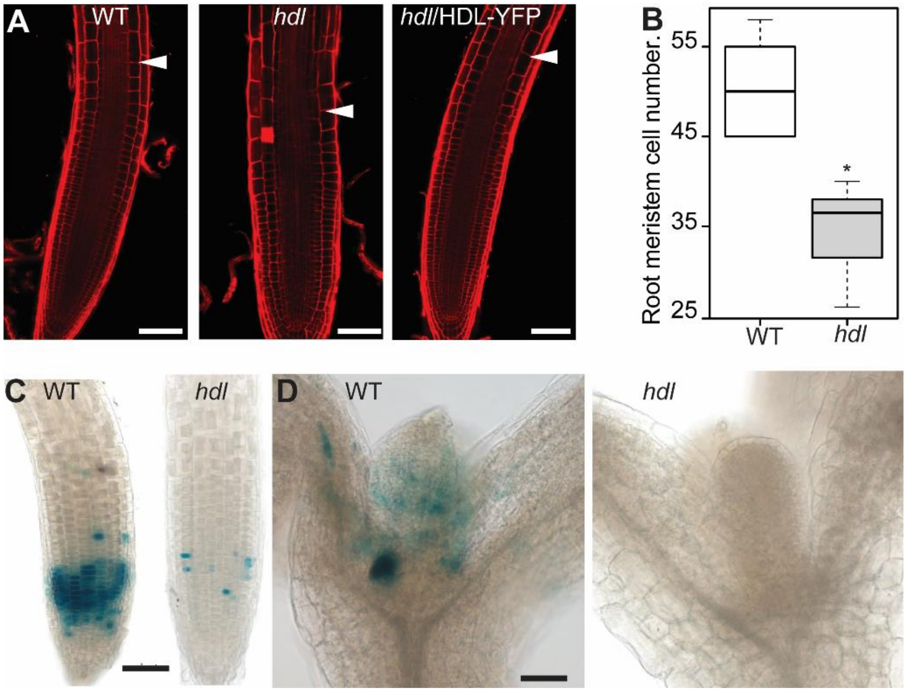


**Figure S4. Cell proliferation is impaired in *hdl* mutant. (A)** Confocal image of 7-dpg root apical meristem of WT, *hdl,* and complementation line. The cell outline was visualized by propidium iodide staining. Arrows indicate the junction between the meristematic and differentiation zones. **(B)** Quantification of root meristem cell number in 10-dpg WT or *hdl,* cells were quantified from the quiescent center (QC) to the first cell of the elongation zone using ImageJ, n=10 seedlings. *, P< 0.05, by Mann-Whitney non-parametric test. **(C-D)** *CYCB1p:GUS* expression in the root apical meristem and shoot apical meristem of 5-dpg WT and *hdl* plants. Scale bar, 50 µm in A,C,D.

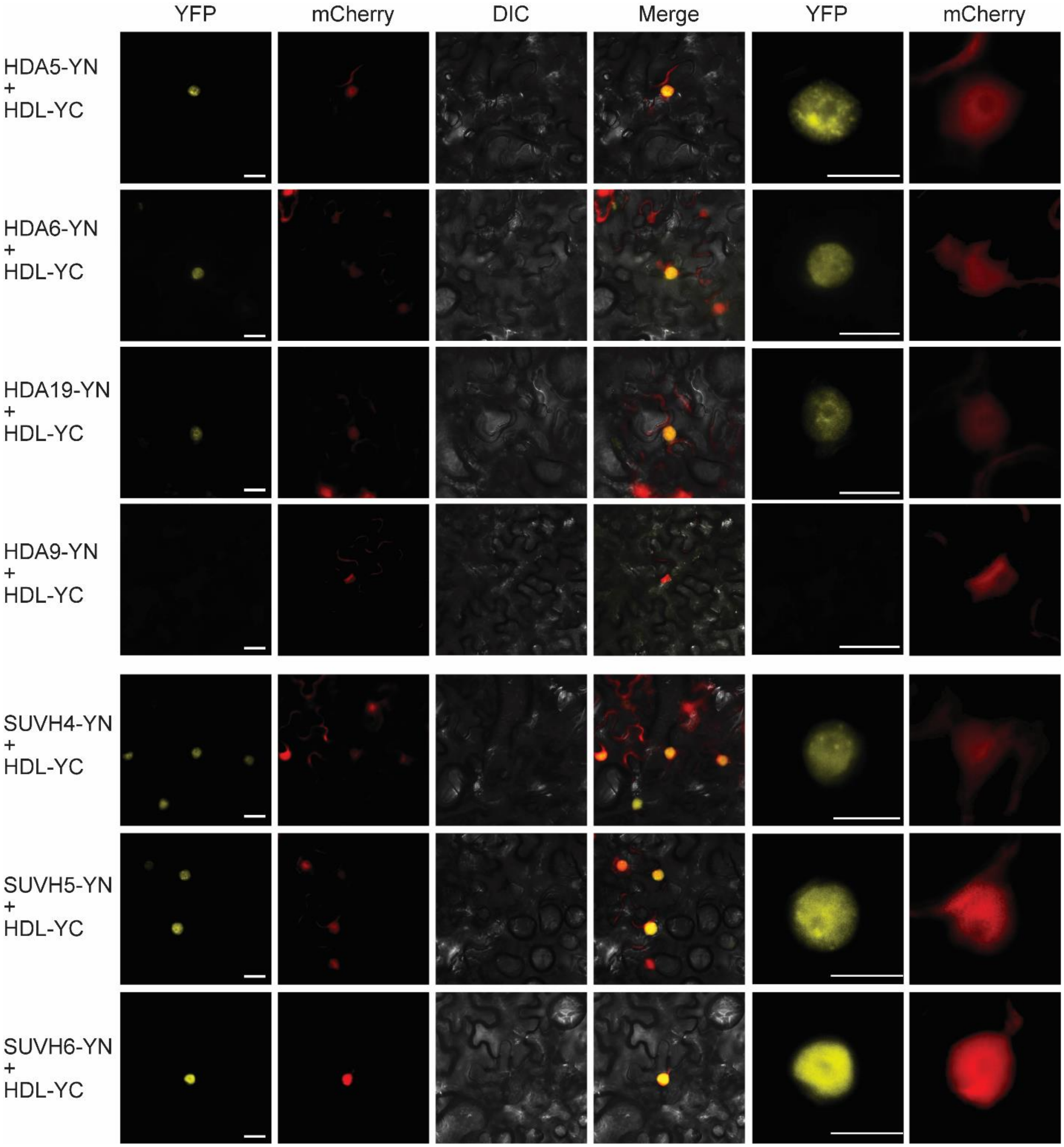


**Figure S5. HDL interacts with HDA5, HDA6, HDA19, HDA19, SUVH4/KYP, SUVH5 and SUVH6. (A)** BiFC assay of HDL interaction with HDA5, HDA6, HDA9, HDA19, SUVH4, SUVH5 and SUVH6. Full-length HDL was cloned with C-terminal YFP in *pEarlyGate202* (YC), and HDA5, HDA6, HDA9, HDA19, SUVH4, SUVH5 and SUVH6 were fused to N-terminal YFP in *pEarlyGate201* (YN). The constructs were co-transformed into abaxial epidermis in *N. benthamiana* using *Agrobacterium* GV3101. The nucleus was visualized with a nuclear marker of mCherry fused with a nuclear localization signal Scale bar 20 μm. The right side of the panel shows the enlarged signal of YFP and mCherry. Scale bar 10 μm.

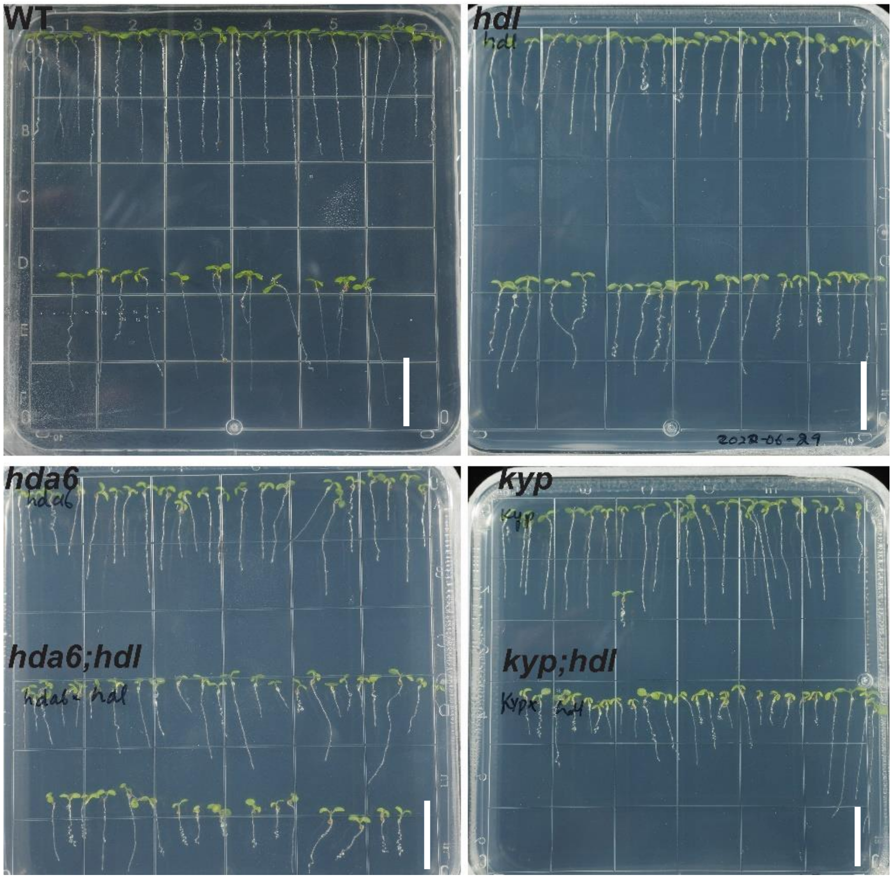


**Figure S6. HDL mutation causes additive effects on *had6* and *kyp*.** Root length phenotype of 6-dpg WT, *hdl, hda6 and hdl,kyp* double mutants. The plants were grown for four days on ½ MS and then transferred to new ½ MS plates for an additional two days. Scale bar, 2 cm.

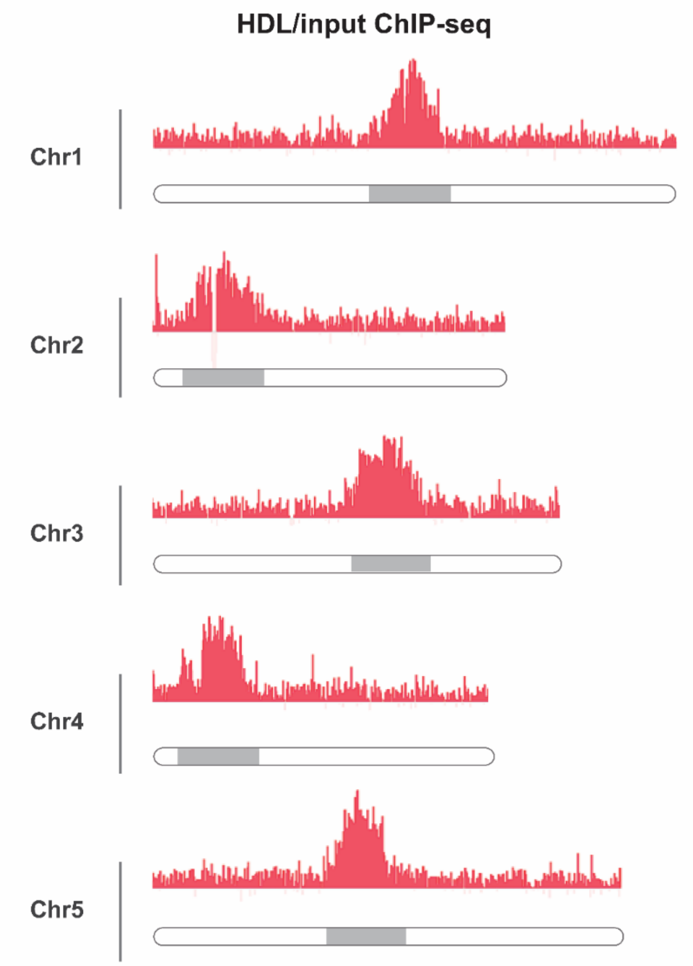


**Figure S7. HDL is enriched in the heterochromatin.** The genome-wide binding profile of HDL ChIP-seq reads across all five chromosomes relative to input DNA. The grey bar at the bottom indicates centromeric and pericentromeric heterochromatin regions.

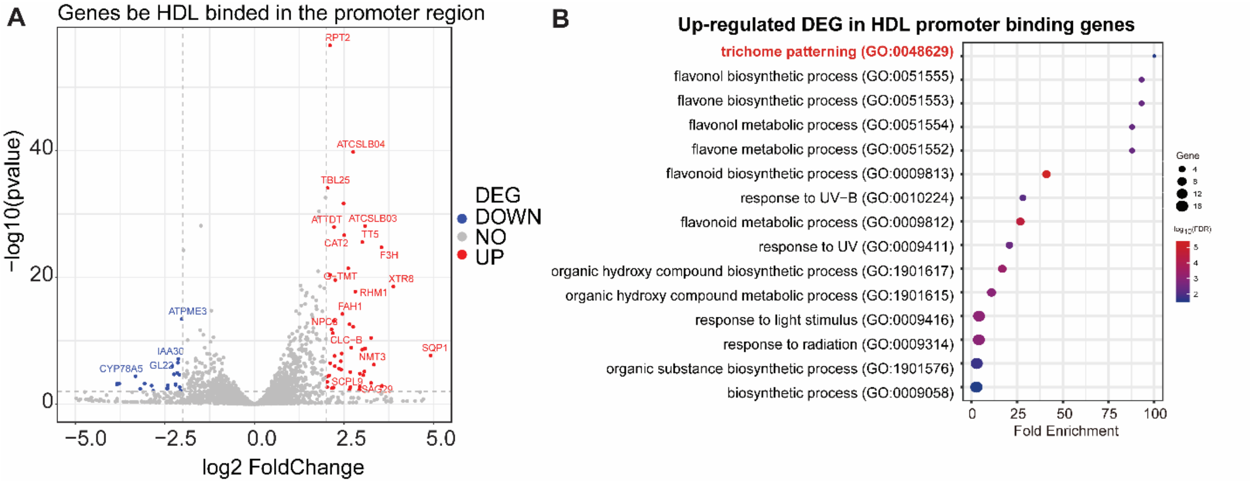


**Figure S8. Differential expression analysis of genes associated with HDL and their associated GO terms. (A)** Transcriptional profile of genes with HDL-associated promoters. Red dots represent up-regulated genes with a log2Fold Change greater than 2 (*hdl*/WT) and a p-value less than 0.01. Blue dots represent down-regulated genes with a log2Fold Change less than -2 and a p-value less than 0.01 in (*hdl*/WT). **(B)** The top enriched GO terms associate with up-regulated genes included trichome patterning and the flavone biosynthesis process. Dot size represents the number of genes, and the color represents the log10 (FDR).

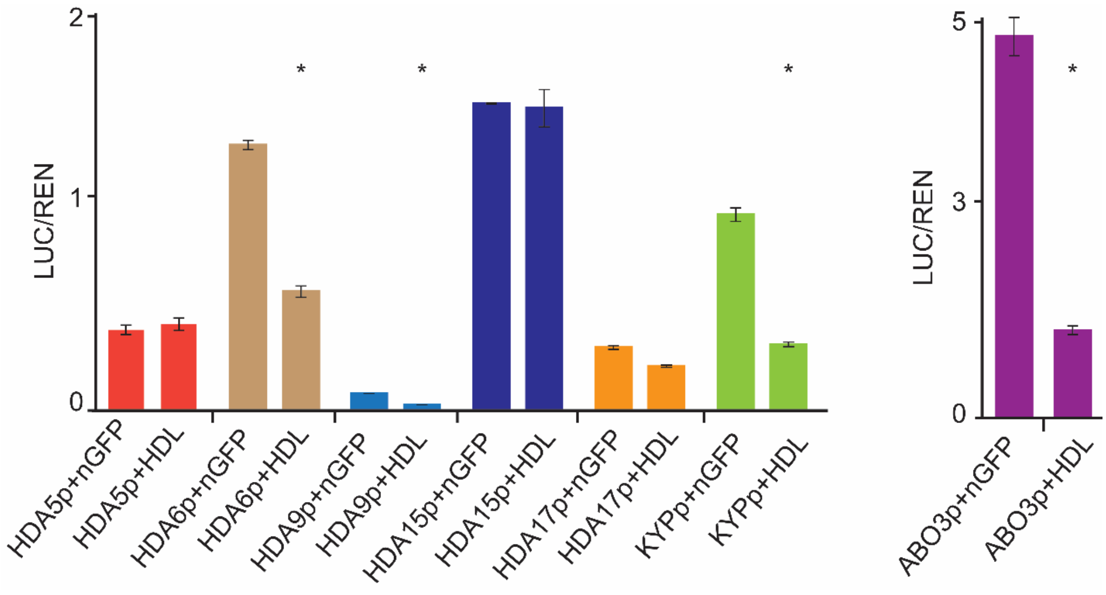


**Figure S9. HDL negatively regulates transcription of HDA6, HDA9, KYP, and ABO3.** Luciferase reporter assay using HDL (35S:HDL) and nucGFP (35S:nucGFP) as effectors, and HDAs, KYP, and ABO3 promoters as reporters. Luciferase activities (LUC) were normalized with the internal control Renilla luciferase (REN). Three independent experiments were performed with two biological samples per experiment. *, $p<0.01$ by Mann whitney non-parametric test.

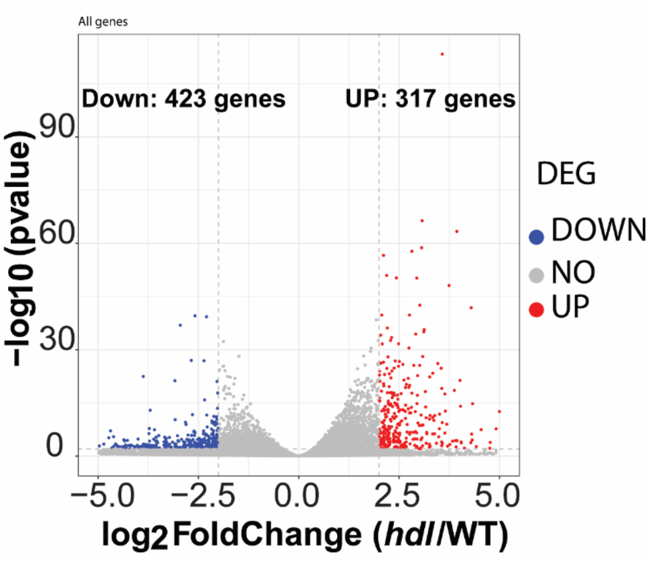


**Figure S10. RNA-seq analysis of WT and *hdl* mutant seedlings.** A volcano plot was used to visualize differentially expressed genes between *hdl* and WT. The comparison was made using the DESeq2 (*20*). A total of 423 downregulated genes (red) and 317 up-regulated (blue) genes were identified with a log2Fold Change greater than 1.5 (*hdl*/WT) and a p-value less than 0.05.

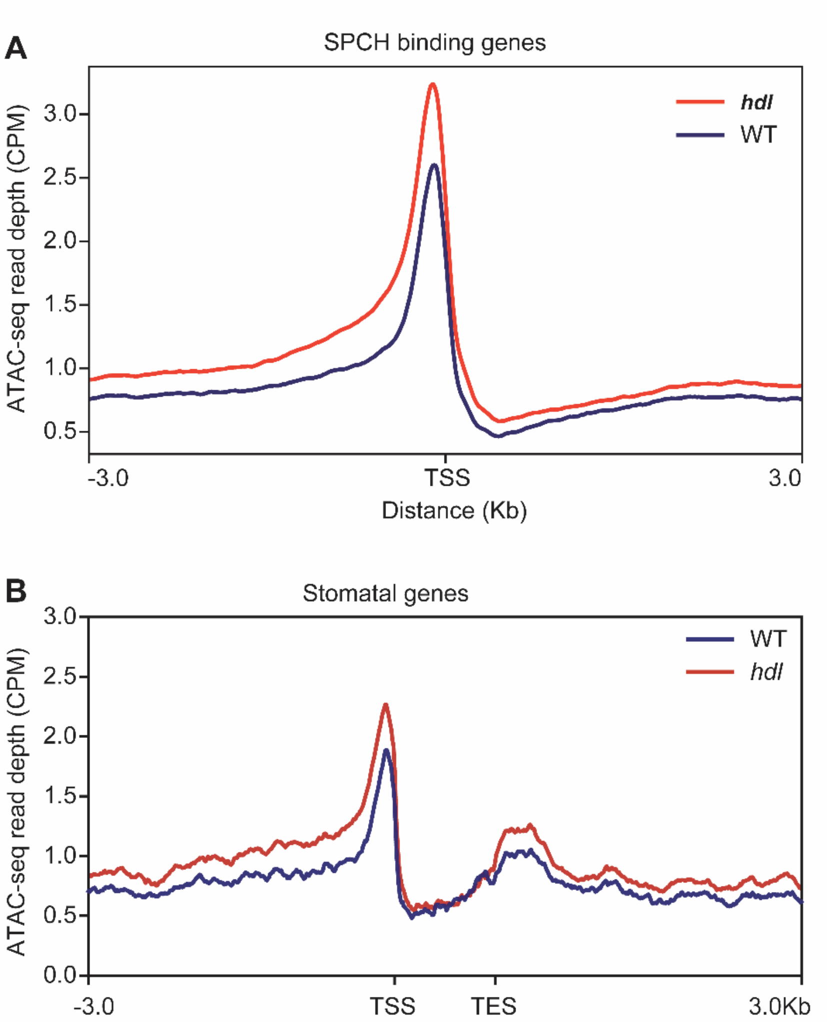


**Figure S11. Increased Chromatin accessibility in SPCH binding targets and stomata-related genes. (A)** ATAC-seq binding profile of SPCH binding genes (*21*) in WT and *hdl*, with the transcription start site (TSS) indicated. **(B)** Average ATAC-seq (CPM) binding profile of all stomata-associated genes (Data S5). The ATAC-seq (CPM) profile upstream of the TSS demonstrated higher chromatin accessibility in *hdl* than WT.

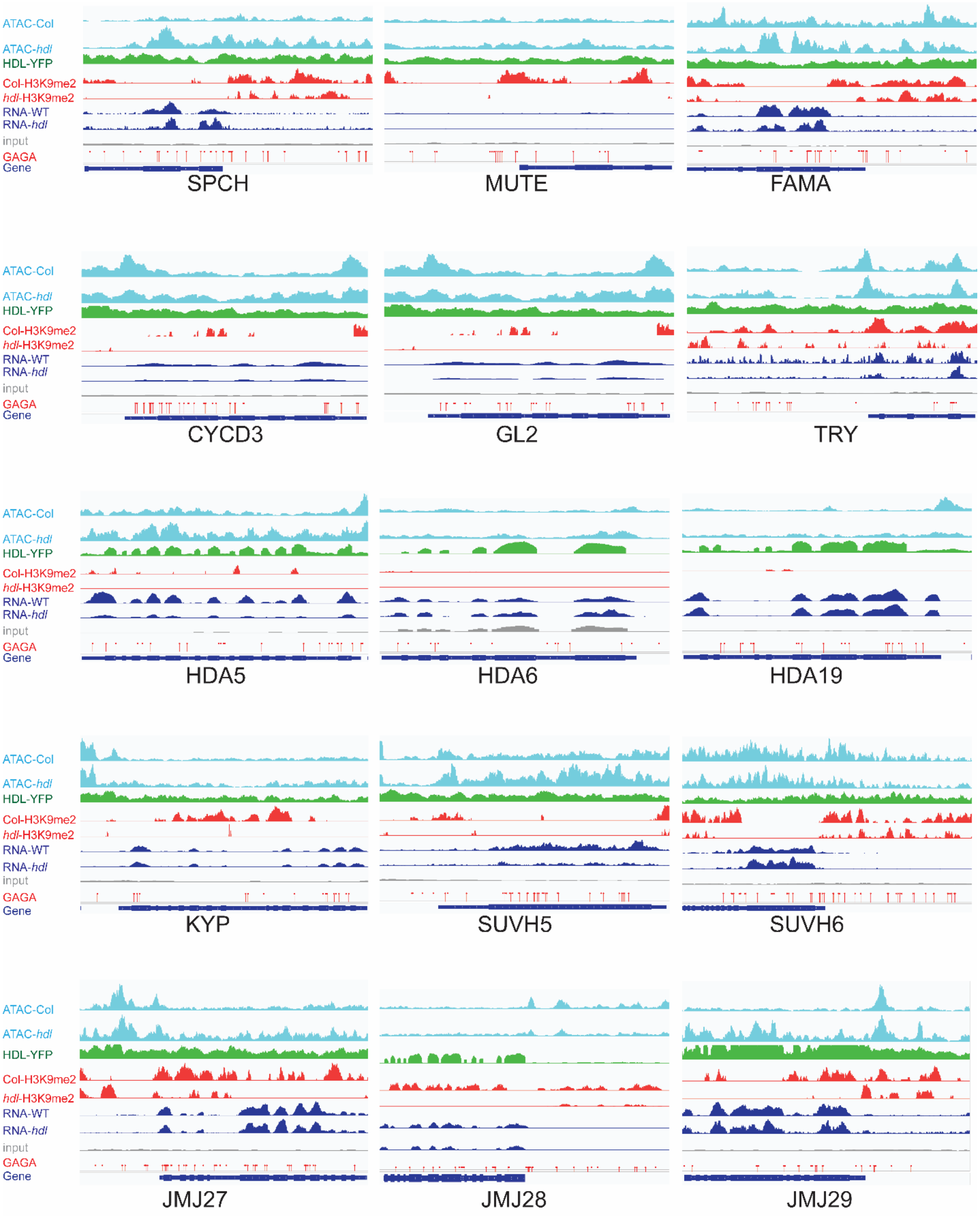


**Figure S12. HDL chromatin association changes the accessibility of cell fate and epigenetic regulators genes.**

A genome browser view of showing chromatin accessibility, HDL binding, H3K9me2 level, transcription in WT, hdl, and GAGA binding motif (red deggers) of stomata, trichome, and chromatin-associated genes.

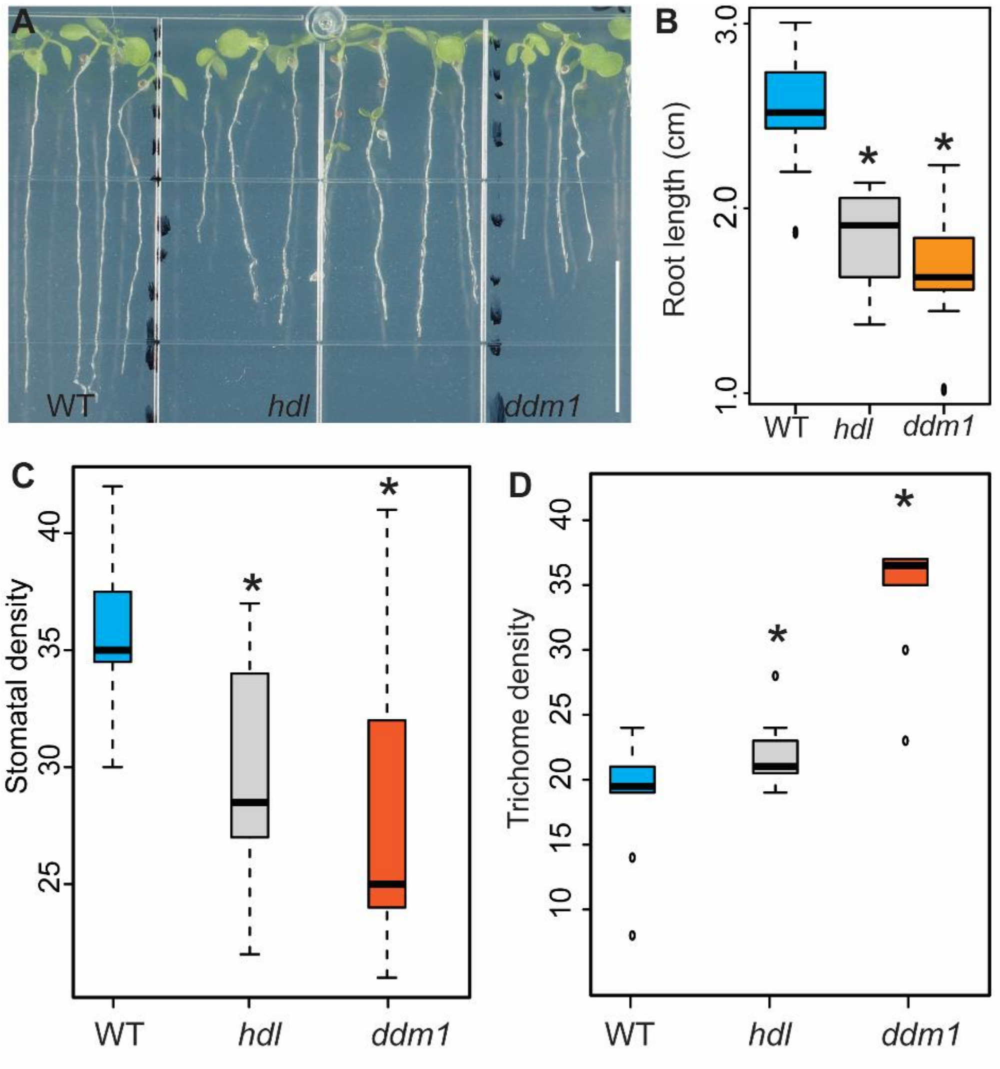


**Figure S13.** ***ddm1*****phenocopies** ***hdl*** **. (A) root phenotypes of WT, hdl and ddm1. (B-C)** Root length and stomata density was decreased in *hdl*, *ddm1* compared to WT. n=18, 14, 24 for WT, *hdl*, and *ddm1* respectively, scale bar 2 mm. **(D)** Trichome density was increased slightly in *hdl*, while *ddm1* showed a more drastic phenotype compared to WT, and *hdl*. A total of 10 adaxial epidermis samples were observed. *, $p < 0.001$, by Mann-Whitney non-parametric test.

**<u>Table S1</u>**
**Primers used in this study**

| **qRT-PCR** | **Sequences 5">3"** |
|---|---|
| SPCH-qRT-PCR-F | ATCATAGGAGGAGTTGTGGAG |
| SPCH-qRT-PCR-R | TAGAACAGGCGGTGAAGGAC |
| MUTE-qRT-PCR-F | CGATCATCGGAGGAGTGATAGA |
| MUTE-qRT-PCR-R | AAGGGAAAGATGGTCGGTTTAG |
| GL2-qRT-PCR-F | ATGAAGCTCGTCGGCATGAGTGGG |
| GL2-qRT-PCR-R | TGGATTGCCACTGAGTTGCCTCTG |
| CPC-qRT-PCR-F | TTGGCGACAGGTGGGAGTTGAT |
| CPC-qRT-PCR-R | AACGACGCCGTGTTTCATAAG |
| TRY-qRT-PCR-F | GTGAGCAGTATCGAATGGGAGT |
| TRY-qRT-PCR-R | CACCGACAAGTCTGTACATTCG |
| TSI-qRT-F | GAACTCATGGATACCCTAAAATAC |
| TSI-qRT-R | GGGAATGGTATCAGATCCTAAC |
| Ta3-qRT-F | GATTCTTACTGTAAAGAACATGGCATTGAGAGA |
| Ta3-qRT-R | TCCAAATTTCCTGAGGTGCTTGTAACC |
| CACTA-qRT-F | GGCTAGCTGTCCGACTCAATGACCT |
| CACTA-qRT-R | CAGACATCCTTTCCTTCAGCTTAGC |
| **Cloning** | |
| HDL-CDS-For | GGGGACAGCTTTCTTGTACAAAGTGGCGATGACGA GAAGAAAATCCACCG |
| HDL-CDS-Rev | GGGGACAACTTTGTATAATAAAGTTGCTTAATCTT CTTCGTCCGTTTCGAC |
| HDL-Pro-F | GCGGCCGCGGCTCTGTGTTTTGGAGTTTCGC |
| HDL-Pro-R | GGCGCGCCTTCAACGAACTGGGGAAAAAC |
| HDL-gRNA-F1 | ATTGGACAAAGGAGTTGAATCTCG |
| HDL-gRNA-R1 | AAACCGAGATTCAACTCCTTTGTC |
| HDL-gRNA-F2 | AAACAAGACTGCATCTACTTCATC |
| HDL-gRNA-R2 | ATTGGATGAAGTAGATGCAGTCTT |
| H2A.W dtopo-Not1-F | caccGCGGCCGCCATGGAATCCACCGGAAAAGTG |
| H2A.W dtopo-Asc1-R | GTCGGCGCGCCCAGCTTTCTTTGGAGACTTGAC |
| H2AW.6pro-Not1-F | GCGGCCGCCGTTAATGTTTCCAATAAC |
| H2AW.6pro-Asc1-R | GGCGCGCCTGCTACGGTTATCGATTAC |
| HDA6-pro-F-Luc | CACCGCGGCCGCAGAGAATCCCATCATTTTGATTC |
| HDA6-pro-R-Luc | TGGCGCGCCCTCCGTCTCTCACTCAGAATC |
| ABO3-pro-F-Luc | CACCGCGGCCGCTGACCCTTTTAGAGACATATCAC |
| ABO3-pro-R-Luc | TGGCGCGCCCTGGTTTAGATTCTGAAATCTTG |
| HDA5-Not1-F | TTAGCGGCCGCGAGATGCGACGGCCTTATTG |
| HDA5-Asc1-R | AGGCGCGCCGCTTGAGGCCGTAGCAGAAGA |

| | |
|---|---|
| HDA9-Not1-F | CACCGCGGCCGCCTTAAAGAAGAAAAGACACG |
| HDA9-Asc1-R | TAGGCGCGCCCATCATTTCTCTCAACATTG |
| HDA15-Not1-F | CACCGCGGCCGCGTCAACACACATTTTTC |
| HDA15-Asc1-R | TAGGCGCGCCGTTACCTCAGTAGTCACTC |
| HDA17-Not1-F | CACCGCGGCCGCGCCTTCAATATATCCTCTG |
| HDA17-Asc1-R | TAGGCGCGCCCAAAACCCGATTCGCAAAG |
| KYP-Not1-F | TATGCGGCCGCCTGGAGACAAGACGACCG |
| KYP-Asc1-F | TTGGCGCGCCGATCACTCTTTTTCCCCTG |
| SPCHp-LUCGib-F | GTAAAACGACGGCCAGTGCCAGTGACGACGATCA CACATAC |
| SPCHp-LUCGib-R | ATGTTTTTGGCATCTTCCATCTCATTTATGTTTTAG ATATAAATA |
| SPCH-F_caccNotI | CACCGCGGCCGCATGCAGGAGATAATACCGGATTT TCTTG |
| SPCH_R_AscI | CGGGGCGCGCCCGCAGAATGTTTGCTGAATTTGTT GAGCC |
| GL2pro-Not1-F | TGCGGCCGCCGGGTAATAGATTAATGGTG |
| GL2pro-Asc1-R | TGGCGCGCCGAATTATGGAAAATATATC |
| GL2-CDS-Not1-F | TGCGGCCGCATGAAGTCGATCGATGGC |
| GL2-CDS-Asc1-R | TGGCGCGCCGCAATCTTCGATTTGTAG |
| **Genotyping** | |
| HDA6-axe-1 | CCCAAAGATATGGAGAGGATAAAG |
| HDA6-axe-2 | CCCAAAGATATGGAGAGGATAAGA |
| HDA6-geno-WA-2 | TAAGACGATGGAGGATTCACG |
| SUVH4/KYP-F | GTACCGACTGAAACGATTGGA |
| SUVH4/KYP-R | AGTTCGGTTGACACATTTTGG |
| LBb1.3 | ATTTTGCCGATTTCGGAAC |
| SALK_052138 LP | TTTCCACTGAAGGAGGAAATTC |
| SALK_052138 RP | TTATTTGACGGTGAAGCCAAG |
| HDL-RT-PCRc-F | GAGGATGGACAGAGGAGTTG |
| HDL-RT-PCRc-R | GCTAACGAGAGTTGTTGGATC |
| HDL-CRISPR-gRNA-For | GAAGATGAGGAACGTATTAAGTG |
| HDL-CRISPR-gRNA-Rev | CCTTTTCCTCTCAGTAAGGC |
| DDM1-geno-F | ATTTGCTGATGACCAGGTCCT |
| DDM1-geno-R | CATAAACCAATCTCATGAGGC |
| HDA6-BiFC-F | GGGGACAAGTTTGTACAAAAAAGCAGGCTA ATGGAGGCAGACGAAATTC |
| HDA6-BiFC-R | GGGGACAACTTTGTATAGAAAAGTTGGGTTT AAGACGATGGAGGATTCAC |
